# Integrative transcriptomics and electrophysiological profiling of hiPSC-derived neurons identifies novel druggable pathways in Koolen-de Vries Syndrome

**DOI:** 10.1101/2024.08.29.610281

**Authors:** A.H.A. Verboven, S. Puvogel, N. Kolsters, B. Latour, K. Linda, E.I. Lewerissa, R. Mijdam, E. Dyke, D. Duineveld, M. Zoodsma, W.J.J. Claassen, A. Oudakker, C. Schoenmaker, D.A. Koolen, B.B.A. de Vries, P.A.C. ’t Hoen, N. Nadif Kasri

## Abstract

Koolen-de Vries Syndrome (KdVS) is a neurodevelopmental disorder (NDD) with no treatment options due to a lack of understanding of its underlying pathophysiology. To investigate neuronal activity in KdVS, human induced pluripotent stem cell (hiPSC)-derived neurons from KdVS and control subjects were cultured on microelectrode arrays (MEAs). Our study identified reduced network burst rates, indicating disorganized network activity in KdVS neurons. To bridge molecular and functional aspects of the syndrome, we developed an experimental framework, MEA-seq, that integrates network activity measurements with high-throughput transcriptome profiling. This approach identified a negative correlation between the expression of the NDD-associated gene *CLCN4* and the network burst rate. Consequently, knockdown of *CLCN4* in KdVS neurons restored the activity to control level, confirming a causal relationship between increased *CLCN4* expression and reduced network burst rate. Additionally, we identified a positive correlation between mitochondrial gene expression and the network burst rate, and identified impaired mitochondrial function in KdVS hiPSC-derived neurons. The transcriptomic signature of KdVS neurons was then used for computational screening against drug perturbation signatures of the LINCS Consortium database, predicting other drug targets and compounds capable of reversing the expression of affected genes in KdVS neurons. We selected 10 compounds for experimental validation, identifying the antioxidant phloretin and the Rho-kinase inhibitor fasudil as potential candidates for restoring the network activity dysfunction in KdVS. We conclude that the integrative molecular and electrophysiological of hiPSC-derived neurons with MEA-seq has excellent potential for identifying novel drugs and druggable pathways for KdVS and other NDDs.

## Introduction

Koolen-de Vries Syndrome (KdVS) is a neurodevelopmental disorder characterized by developmental delay, intellectual disability, hypotonia, dysmorphic facial features, and increased social behavior [1]. KdVS is caused by haploinsufficiency of the lysine transferase 8 regulatory non-specific lethal complex unit 1 (KANSL1), which can originate from a heterozygous variant in *KANSL1* [2, 3] or, more commonly from a heterozygous microdeletion in 17q21.31 that includes *KANSL1* [4, 5] and is flanked by extensive low copy repeats or segmental duplications. Despite knowing the underlying genetic diagnosis there is little insight in the pathological mechanisms and therefore, few treatment options are available. Since neuronal network functioning may be compromised in KdVS [6, 7], understanding the molecular changes leading to human neuronal activity heterogeneity may help to identify potential new targets for treating KdVS, as well as other neurodevelopmental disorders.

KANSL1 functions as a scaffolding protein by binding other subunits of the non-specific lethal complex (NSL), including the acetylase lysine transferase 8 (KAT8) [8]. KAT8 is an epigenetic regulator responsible for gene activation via H4K5 and H4K8 acetylation when bound to the NSL complex, whereas it mainly acetylates H4K16 as part of the male-specific lethal (MSL) complex [9]. The function of KANSL1 during development has been studied *in vivo* using mutant *Drosophila* and mouse models [2, 7], showing reduced learning abilities and impaired recognition and associative memory linked to *KANSL1* deficiency. Gene expression changes pointed towards impaired synapse function and neurogenesis associated with KANSL1 haploinsufficiency in mice [7]. Additionally, an *in vitro* study of KdVS-derived fibroblasts showed imbalanced cellular and metabolic homeostasis [10], which is in line with the known functions of KANSL1 and the NSL complex [11–14]. Using human induced pluripotent stem cell (hiPSC)-derived neurons, we have previously shown oxidative stress-induced autophagosome accumulation at excitatory synapses of KdVS hiPSC-derived neurons, in combination with reduced synaptic puncta and impaired neuronal network activity [6]. Together, these results indicate altered neuronal function in KdVS. However, the connection between molecular changes and neuronal functioning in KdVS is not yet understood, which precludes the development of effective treatment for the disorder.

hiPSCs enable the modeling and study of different stages of human development in health and disease. Transcriptome analysis (RNA-seq) can be used to gain insight into the molecular mechanisms underlying these developmental changes; therefore, it has been widely used for the study of neurodevelopmental disorders [15–17]. The identification of transcriptional signatures associated with disease-related neuronal function can be used to repurpose drugs that could counteract relevant and specific transcriptional changes, and thus, may rescue the neuronal activity phenotypes. For instance, the NIH LINCS L1000 project is a database consisting of gene expression profiles of more than 30,000 genetic and drug perturbations; representing a potentially ideal platform to identify compounds for drug repurposing [18, 19]. However, RNA-seq experiments are often performed separately from functional experiments and it is unclear which changes in gene expression are directly related to neuronal phenotypes. To fill this gap, Bardy et al. designed Patch-seq, which combines morphological, electrophysiological and gene expression measurements in single human neurons [20]. However, neurons communicate and function within a network, resulting in activity patterns that are missed when studied at single cell level.

Microelectrode arrays (MEAs) are culture plates with embedded electrodes that facilitate recording of neuronal network activity in a semi high-throughput manner. We previously showed that *Ngn2*-induced hiPSC-derived neurons grown in MEAs exhibit a robust and consistent neuronal network phenotype across different control cell lines [21], and allow for disease-specific phenotyping [6, 21–23]. Here, we present MEA-seq, a novel approach that allows for integration of gene expression and neuronal network activity measurements using microelectrode arrays in the same experiment. We used this approach to study the mechanisms underlying neuronal network dysfunction in KdVS hiPSC-derived neurons. In addition, we implemented a computational approach for drug identification for KdVS, based on the transcriptomic signatures of the LINCS database. Subsequently, we studied the effect of selected compounds on neuronal network functioning *in vitro*. Overall, this study yields new insight into the neuronal mechanisms underlying KdVS by integrating information at both molecular and functional level and presents a new framework for the development and testing of treatments for neurodevelopmental disorders.

## Materials and Methods

### hiPSCs lines

In this study we used three control hiPSCs, four KdVS patient derived lines, and two CRISPR/Cas9 edited hiPSC lines. Two of the control lines were commercially available hiPSCs. C1 is wtc11 (UCSFi001-A) derived from a 30-year old male [24], and C2 is from a 36-year old female [25]. We generated a third hiPSCs control line (C3) from fibroblasts of the healthy mother of one of the KdVS individuals included in the study (KdVS1). Fibroblasts were obtained from the four KdVS individuals, presenting the full spectrum of KdVS associated symptoms. KdVS1 [26] and KdVS2 cell lines [3, 27] derived from female KdVS patients with a heterozygous mutation in exon 2 of *KANSL1* that leads to a premature stop codon. The KdVS3 cell line originates from a female patient with a heterozygous 17q21.31 microdeletion [26]. The KdVS4 cell line derives from a female patient with a heterozygous mutation in intron 12 of *KANSL1*, predicted to result in a frameshift and premature stop codon due to loss of exon 13 (**Fig. S1A-B**) [26]. Reprogramming procedure is detailed in Supplementary methods. hiPSCs clones from C3 and KdVS1-4 have been previously validated through quality control tests including morphological assessment and karyotyping to confirm genetic integrity. All selected hiPSCs clones from the different lines were positive for pluripotency markers (OCT4, TRA-1-81, and SSEA4) (**Fig. S1C**) [6, 24]

### CRISPR/Cas9 hiPSCs line generation

The genetically edited CRISPR/Cas9 iPSC line *KANSL1*-CRISPR is the line originally described in Linda et al. 2022 [6], with a heterozygous nonsense mutation in exon 2 of *KANSL1*. A CRISPR/Cas9 genetically edited line derived from C1 was created via the introduction of a heterozygous nonsense mutation in exon 2 of *KANSL1* (NC_000017.10(NM_015443.4): c.428_429insN; *KANSL1*-CRISPR-C1) using the sgRNA CTTAGAACCATGAATACGAG. *KANSL1*-CRISPR-C1 was found to be karyotypically normal. This line was used in the second round of drug experiments to validate our findings in an isogenic pair.

### Generation of rtTA/Ngn2+ hiPSCs

rtTA/Ngn2-positive hiPSCs were generated for all lines, using lentiviral vector to stably integrate the transgene into the genome, as described previously [6]. hiPSCs were grown on 6-well plates coated with Matrigel (Corning, #356237) and cultured on E8 Flex medium (Thermo Fisher Scientific), supplemented with primocin (0.1 μg/ml, Invivogen) and low concentration of puromycin (0.5 µg/mL) and G418 (50 μg/ml), at 37°C/5% CO_2_. Medium was refreshed every other day, and cells were passaged using ReLeSR (Stem cell technologies) 1-2 times per week.

### Neuronal differentiation

hiPSCs were differentiated into excitatory cortical neurons by doxycycline-induced overexpression of Neurogenin-2 (Ngn2), as described previously [28]. rtTA/Ngn2-positive hiPSCs were plated on 24-well microelectrode arrays (Multichannel Systems, MCS GmbH, Reutlingen, Germany) or on nitric-acid treated coverslips in 24-well plates. Well and coverslips were previously coated with 50 µg/mL poly-L-ornithine hydrobromide (PLO; Sigma-Aldrich, #P3655-10MG) in borate buffer (50 mM) for at least 3 hours at 37°C/5% CO_2_, followed by coating with 5 µg/mL of human recombinant laminin 521 (BioLamina, #LN521-02) in DPBS (with calcium and magnesium, Gibco #14040117) overnight at 4 °C. Control hiPSCs were plated at ∼20,000 cells/well in E8 basal medium (Gibco, #A1517001) supplemented with 0.1 μg/ml primocin (Invivogen, ant-pm-2), 1% RevitaCell (Thermo Fisher Scientific, #A2644501), and 4 µg/mL doxycycline (Sigma-Aldrich, #D9891-5G). KdVS hiPSCs were plated at a higher density to compensate for cell death and ensure similar cell density after full neuronal differentiation. Differentiation procedure is detailed in Supplementary methods.

### Microelectrode array recordings

Spontaneous activity was recorded in hiPSC-derived neurons from all control and KdVS cell lines, from DIV16 up to a maximum of DIV79, with measurements performed every week. The recordings were performed using the 24-well MEA system (Multichannel Systems, MCS GmbH, Reutlingen, Germany) and 48-well plates of Axion Biosystems (drug test). For MCS recordings, spontaneous neuronal network activity was registered for 10 minutes, after a 10-minute acclimatization period, in a recording chamber that was maintained at 37°C/95% O_2_/5% CO_2_. The signal was sampled at 10 kHz and filtered with a high-pass 2^nd^ order Butterworth filter with a 100 Hz cut-off frequency and a low-pass 4^th^ order Butterworth filter with a 3500 Hz cut-off frequency. The noise threshold for spike detection was set at ± 4.5 standard deviations. Data analysis was performed off-line using Multiwell Analyzer software (Multichannel systems) that permitted extraction of spike-trains. The burst detection was adjusted by setting the minimal interval between bursts to 700 ms. A custom-made MATLAB (The Mathworks, Natick, MA, USA) code was used to extract parameters describing the spontaneous network activity. For recordings using Axion Biosystems, network activity was measured for 5 minutes in a recording chamber maintained at 37°C/5% CO_2_. Data analysis was performed using Axion Biosystems’ NeuralMetricsTool software, followed by offline analysis using a custom-made R script. We calculated the mean firing rate (MFR, spikes/s), the mean percentage of random spikes (PRS, %), the burst rate (BR, bursts/min), the burst duration (BD, in sec), the inter-burst interval (IBI, in sec), the burst spike rate (BSR, spikes/s), the network burst rate (NBR, NB/min), the network burst duration (NBD, in sec), the interval between network bursts (NIBI, in sec), and the coefficient of variation calculated on the NIBI (CV_NIBI_), which reflects the regularity of the NBR. The wells that at DIV30 were showing MFR <0.1 spikes/s and BR < 0.4 bursts/min were excluded from analysis. Linear discriminant analysis (LDA) was performed on five parameters affected in KdVS (PRS, IBI, NBR, NIBI, CV_NIBI_), using the lda function from MASS (R package). Group predictions were determined based on LDA on all 10 MEA parameters.

### RNA sequencing

RNA sequencing was performed in hiPSCs and hiPSC-derived neurons co-cultured with rat astrocytes, from control lines (C1-3), *KANSL1*-CRISPR, and three KdVS lines (KdVS1-3). RNA from hiPSC-derived neurons was isolated after measuring neuronal network activity on MEAs at DIV30, from 3 replicates per line, with the seven lines split into two batches (batch 1 = C3, *KANSL1*-CRISPR, KdVS1; batch 2 = C1-2, KdVS2-3). RNA was isolated with the Quick-RNA Microprep kit (Zymo Research, R1051) according to manufacturer’s instructions. RNA quality was checked using Agilent’s Tapestation system (RNA High Sensitivity ScreenTape and Reagents, 5067-5579/80). RIN values ranged between 8.4 – 9.8 for hiPSCs, and 6.4 – 8.8 for hiPSC-derived neurons. Library preparation was performed using a published single-cell RNA-seq protocol [29], adapted for bulk RNA-seq experiments (Details in Supplementary information). Libraries were sequenced on the NextSeq 500 platform (Illumina) using a V2 75 cycle kit (Read 1: 18 cycles, Read 2: 52 cycles, Index 1: 10 cycles). RNA-seq data pre-processing steps are detailed in Supplementary methods.

### RNA-seq data analysis

Data analysis was performed in R. Counts were normalized with TMM method and transformed to counts per million (cpm) using edgeR v3.26.8 [30]. Only transcripts with cpm>2 in at least four samples for hiPSCs, or at least three samples for hiPSC-derived neurons, were included for further analysis. Counts belonging to the Y chromosome were excluded. For the principal component analysis (PCA) including all cell types, rat counts were converted to human homologues, and counts from all cell types were voom-transformed (log2-transformation on cpm values) using limma v3.40.6. PCA was then performed on voom-transformed data using the *prcomp* function from the stats package in R. Differential expression (DE) analysis between KdVS and control neurons was performed using limma [31]. We corrected for batch effect. *duplicateCorrelation* function was used to account for variability within each cell line. Genes with Benjamini-Hochberg (BH)-corrected p<0.05 were considered as significantly differentially expressed. RNAseq analysis for querying the LINC database are detailed in Supplementary methods.

### MEA-seq correlation analysis

Spearman correlations were calculated between the expression profiles of hiPSC-derived neurons and the 10 MEA parameters describing neuronal network activity at DIV30, using *corr.test* of the psych package v2.0.12. Voom-transformed counts, not corrected for batch effect, were used as input for the correlation analysis. Correlations with a Benjamini-Hochberg (BH)-corrected *p*<0.05 were considered significant. Post-hoc analysis was performed on the significant correlations, using control samples only, KdVS samples, and female samples only, to ensure that correlations were not observed solely by differences between condition or by gender. Correlations between genes and MEA parameters with unadjusted *p*<0.05 in all three post-hoc analysis groups were considered significant. Then, we performed gene set enrichment analysis (GSEA; fgsea v1.10.1 [32]) on the genes ranked according to their correlation with the specific MEA parameter. Gene symbols (Ensembl version 94) were converted to gene symbols of Ensembl version 97, corresponding to the version of gene symbols used in MSigDB.

### CLCN4 RNA interference

For RNA interference (RNAi) knockdown experiments, a DNA fragment encoding a short hairpin RNAs (shRNA) directed against *CLCN4* was cloned into a lentiviral vector by VectorBuilder (pLV[shRNA]-EGFP:T2A:Puro-U6>hCLCN4[shRNA#3]; vector ID: VB900037-1851bkf). The *CLCN4* target sequence was: CCCGAATCCATCATGTTTAAT. A scrambled shRNA was used as non-targeting control, cloned into a lentiviral vector by VectorBuilder (pLV[shRNA]-EGFP:T2A:Puro-U6>Scramble_shRNA#1; vector ID: VB010000-0009mxc) with the following hairpin sequence: CCTAAGGTTAAGTCGCCCTCG. An empty vector only expressing GFP was used as control vector. Lentiviral particles were prepared, concentrated, and tittered as described previously [34] (Details in Supplementary information). For *CLCN4* knockdown experiments, DIV1 KdVS1 hiPSC-derived neurons were transfected with lentivirus expressing shRNA’s or GFP for 4-6 hours, 3 wells per condition.

### Immunocytochemistry

hiPSC-derived neurons (DIV23 or DIV30) were washed with ice-cold 1X PBS (Sigma-Aldrich, P5493-1L) and fixed or with 4% paraformaldehyde / 4% sucrose. Next, cells were washed, permeabilized and incubated with primary antibodies overnight at 4°C, washed again, and incubated with secondary antibodies for 1 hour at room temperature (Detailed steps and information of the antibodies in Supplementary information). Imaging was performed on a Zeiss Axio Imager Z1 at 20X (MAP2/GFAP) or 63X (MAP2/CLCN4) magnification.

### Quantification of mRNA by RT-qPCR

RNA was isolated with Quick-RNA Microprep kit (Zymo Research, R1051), according to manufacturer’s instructions. RNA samples were converted into cDNA by the iScript cDNA synthesis kit (BIO-RAD, 1708891). cDNA products were clean up using the Nucleospin Gel and PCR clean-up kit (Macherey-Nagel, 740609.250). Primers targeting *CLCN4* were designed using the NCBI Primer-BLAST tool [35]. RT-qPCR was performed using the Quantstudio 3 apparatus (Thermo Fisher Scientific) with GoTaq qPCR master mix 2X with SYBR Green (Progema, A600A), according to the manufacturer’s protocol (primers and RT-qPCR detailed in Supplementary information). All samples were included in triplicate. The arithmetic mean of the Ct values of the technical replicas was used for calculations. Relative mRNA expression levels were calculated using the 2^-ΔΔCt^ method and normalized to the expression of the housekeeping gene *GADPH*.

### Seahorse Mito Stress Test

To avoid a possible buffering of mitochondrial function by wild-type astrocytes, this experiment was performed in pure neuronal cultures at DIV7. Oxygen consumption rates (OCR) were measured using the Seahorse XFe96 Extracellular Flux analyzer (Seahorse Bioscience) in control (C1, C2; n_wells_=23-24) and KdVS (KdVS1, KdVS3; n_wells_=24) hiPSC-derived neurons. At DIV0, control hiPSCs were plated at a density of ∼10,000 cells/well. KdVS hiPSCs were plated at a higher density to compensate for cell death and ensure similar cell density after full neuronal differentiation. We added rat astrocyte-48 hours conditioned Neurobasal medium (Gibco, #21103-049) supplemented with 20 μg/mL B-27 (Gibco, #0080085SA), 1% GlutaMAX (Gibco, #35050038), 0.1 µg/mL primocin, 4 µg/mL doxycycline, 10 ng/mL human recombinant NT-3, and 10 ng/mL human recombinant BDNF. Then, neuronal differentiation was performed as described above, up to DIV7.

At DIV7, one hour before measuring, culture medium was replaced by Agilent Seahorse XF Base Medium (Agilent #103334-100) supplemented with 10 mM glucose (Sigma), 1 mM sodium pyruvate (Sigma), and 200 mM L-glutamine (Gibco) and incubated at 37°C without CO2. Basal oxygen consumption was measured three times followed by three measurements after each addition of 2 µM of oligomycin A (Sigma #75351-gMG), 1-4 µM carbonyl cyanide 4-(trifluoromethoxy) phenylhydrazone (FCCP; Sigma #C2920-50MG), and 0.5 mM of rotenone (Sigma #R8875-25G) and 0.5 mM of antimycin A (Sigma #A8674-100MG), respectively. One measuring cycle consisted of 3 minutes of mixing, 3 minutes of waiting and 3 minutes of measuring. The oxygen consumption rate (OCR) was normalized to citrate synthase (CS) activity, to correct for the mitochondrial content of the samples [36]. The CS activity was measured according to [37], modified for Seahorse 96-wells plates. We recorded background CS activity in the absence of its substrate oxaloacetic acid, followed by adding oxaloacetic acid, and subtracting the initial activity from the activity in the presence of substrate. After measuring the OCR, the Seahorse medium was replaced by 0.33% Triton X-100 in 10 mM Tris-HCl (pH 7.6). Next, the plates were stored at -80°C. Before measurements, the plates were thawed and 3 mM acetyl-CoA (Sigma), 1 mM DTNB (Sigma), and 10% Triton X-100 were added. The background conversion of DTNB was measured spectrophotometrically at 412 nm and 37°C for 10 minutes at 1-minute intervals, using a Tecan Spark spectrophotometer. Subsequently, the reaction was started by adding 10 mM of the CS substrate oxaloacetate (Sigma), after which the ΔA_412_ nm was measured again for 10 minutes at 1-minute intervals. The CS activity was calculated from the rate of DTNB conversion in the presence of oxaloacetate, subtracted by the background DTNB conversion rate, using an extinction coefficient of 0.0136 mmol^-1^.cm^-1^.

OCR levels normalized to CS activity were used to calculate the parameters describing mitochondrial function. For the FCCP condition, we chose the concentration that resulted in the highest OCR levels, for each cell line independently (C1 and KdVS1 = 4 µM; C2 and KdVS3 = 3 µM). The basal respiration was calculated by subtracting the non-mitochondrial respiration rate (measurements after rotenone and antimycin A treatment) from the last basal respiration rate measured before oligomycin treatment. The maximal respiration was calculated by subtracting the non-mitochondrial respiration rate from the maximum rate measured after FCCP injection. The ATP production was calculated by subtracting the minimum rate measured after oligomycin treatment from the last measured rate before oligomycin treatment. Extracellular acidification rate (ECAR) was also normalized to CS activity and the average normalized ECAR was calculated for the first three basal respiration measurements.

### Querying the LINCS database

Query genes were selected per individual KdVS line (detailed in Supplementary methods). First, the DE gene lists were filtered for best inferred genes (BING) from the LINCS database, consisting of 10,174 genes including 978 landmark genes and 9,196 computationally well-inferred genes [19]. In total, 5,885 out of 8,817 DE genes remained for further analysis. The top 150 up-regulated and 150 down-regulated genes, sorted by the t-statistic, were selected as query genes per KdVS line. LINCS Phase I (GSE82742) and Phase II (GSE70138) level 5 data, collectively accounting for ∼600,000 gene expression profiles, were used to query the KdVS genes. These include gene expression profiles for 30,780 perturbagens, profiled at different concentrations, different times of exposure, and across different cell types. The level 5 data were stored internally using MongoDB v4.2.6. We calculated the connectivity score as previously published [19] (detailed in Supplementary information), to assess the extent of connectivity between our biological state of interest (i.e. KdVS query genes) and the gene expression profiles in the LINCS database (ordered list of z-scores for 10,174 BING). Weighted connectivity scores were then normalized to account for differences between different cell lines and perturbation types. Finally, we summarized the connectivity score of each perturbagen across all cell lines on which this specific perturbagen was profiled, generating “cell-summarized” quantile scores.

### Compound selection

We selected compounds for validation on a MEA screen via two approaches: **1)** We calculated the overlap in the mechanisms of action (MOA) of the compounds with negative connectivity scores with all KdVS lines (n=193), and with the *KANSL1* mutation lines (n=536). Among compounds that were in the clinical phase “launched”, as specified in the clue drug repurposing tool [38], we selected one compound per top MOA category. These included FDA-approved compounds (drugs@FDA, version 27-10-2020), among others. In addition, we only selected compounds with targets expressed in our hiPSC-derived neurons (cpm > 10). **2)** We selected the compounds with the largest absolute negative quantile scores (Q_c,t_): overlapping compounds between KdVS lines with a Q_c,t_ < -0.9 (n=55), and between *KANSL1* mutation lines with a Q_c,t_ < -1 (n=80). Then, we only kept launched compounds with targets expressed in our hiPSC-derived neurons. In total, 10 compounds were selected for *in vitro* testing on MEA.

### Drug treatment

*KANSL1*-CRISPR hiPSC-derived neurons were used for drug testing. Romidepsin and (RS)-(±)-sulpiride were dissolved in DMSO to a final concentration of 10 mM. Meclofenamic acid was dissolved in Milli-Q. All compounds were further diluted in Milli-Q to a concentration of 1 mM (in 10% DMSO). Cabazitaxel and ezetimibe were insoluble in Milli-Q; therefore, these were further diluted in DPBS (Gibco, #14040117) with addition of 0.25% Tween-20 (Merck, #8.22184.0050) to a concentration of 1 mM (in 10% DMSO). Compounds (1 mM) were added to the neuronal cultures on MEAs, from DIV6 until DIV44, in a 1:100 ratio to maintain a final concentration of 10 µM (in 0.1% DMSO) during differentiation. Prednisone, fasudil, and phloretin were also tested at concentrations of 1 µM and 0.3 µM in a second round of experiments using C1 and *KANSL1*-CRISPR_C1 neurons. DMSO and DMSO + Tween-vehicle controls were included, by adding DMSO at a ratio of 1:1000 (0.1% final concentration) and by adding Tween-20 (0.25% in neurobasal medium) at a ratio of 1:100 (0.0025% final concentration), in the neurobasal medium.

### Cell viability

Cell death was assessed at DIV44, after the last MEA measurement. The level of cell death was determined using propidium iodide (PI; ThermoFisher Scientific, BMS500PI). PI in DPBS (1:3000) was added to the neurons for 5 min. As a positive control, one well was treated with PI and Triton X-100 (0.2% final concentration). Pictures were taken with the ZOE Fluorescent Cell Imager at 556/615 nm excitation/emission.

### Statistics

To evaluate the effect of the condition (control vs. KdVS) on activity-related variables measured over time and including multiple cell lines per condition, we used linear mixed-effect modelling with *lmer* function of lme4 [39]. We compared the fitting between models with and without condition as a covariable, using *anova* function of stats. If significant (*p*-value < 0.05), an unpaired *t*-test was performed per time point. For multiple comparisons between conditions over time including only one cell line per condition, or multiple comparisons between conditions considering only one measurement, we performed ANOVA using *aov* function of stats, followed by a Tukey honest significance difference (HSD) post-hoc test with *TukeyHSD*. To compare between the conditions at a single time point, an unpaired *t*-test was performed. *p*-values < 0.05 were considered significant. For comparisons between all compound-treated conditions and DMSO-treated *KANSL1*-CRISPR (control), an ANOVA was followed by Dunnett’s multiple comparisons post-hoc test using the *DunnettTest* function of DescTools [40]. Adj. *p*-values < 0.05 were considered significant. The MEA recordings for the second batch of drug experiments were conducted using an Axion Biosystems machine and the NeuralMetricsTool. This approach captures numerous activity-related parameters, enabling a multiparametric characterization of network activity. To evaluate whether the tested compounds induced effects relative to the control phenotype, we performed PCA on normalized and scaled parameters for each drug and DIV using the *prcomp* function from the stats package. We then applied UMAP to the principal components (PCs) that accounted for more than 2% of the variance in the data. The number of PCs considered is indicated in the corresponding figures. Next, we conducted hierarchical clustering analysis based on the two UMAP dimensions, using Euclidean distances. This was done with the *hclust* function from the stats package, with the method set to “complete”. We used *cutree* function with k=2 to obtain two distinct activity clusters. To test whether adding the compound would shift KdVS-treated networks closer to the control phenotype, we evaluated the association between condition (control, KdVS, or compound-treated KdVS) and each activity cluster using a Bayesian generalized linear model. This model assessed the effect of condition on the odds of a network being assigned to each cluster. We used the *stan_glm* function from the rstanarm R package with a binomial distribution and a logit link function. We obtained 90% credible intervals for the estimates (ln(odds); logits) from the posterior distribution. To convert the logits to probabilities displayed in the plots, we exponentiated them using Euler’s number and subsequently calculated the probability as odds/(1+odds). The number of wells or the number of cells (n) included in each analysis is reported in the figure legends.

## Results

### Synchronized neuronal network activity is reduced in hiPSC-derived neurons from KdVS individuals

We generated hiPSC-derived neurons from a cohort of four KdVS individuals (KdVS1-KdVS4) and three controls (C1 and C2 are independent controls, while C3 line is derived from the healthy mother of KdVS1; **Fig. 1A**)[6]. Additionally, we used a previously characterized *CRISPR*/*Cas9* generated isogenic cell line with a gene dose-dependent deficiency of *KANSL1* from C3 (*KANSL1*-CRISPR) [6]. The genomic region affected in individuals with KdVS is also associated with a common 900 kb inversion polymorphism, which is present in individuals with a H2 haplotype, yet absent in individuals with H1 haplotype. We investigated the haplotypes in our cell lines and observed that most lines carry the H1/H1 or H1/H2 haplotype, whereas KdVS3 carried the H2/H2 haplotype (**Fig. S1D**).

**Figure 1.**
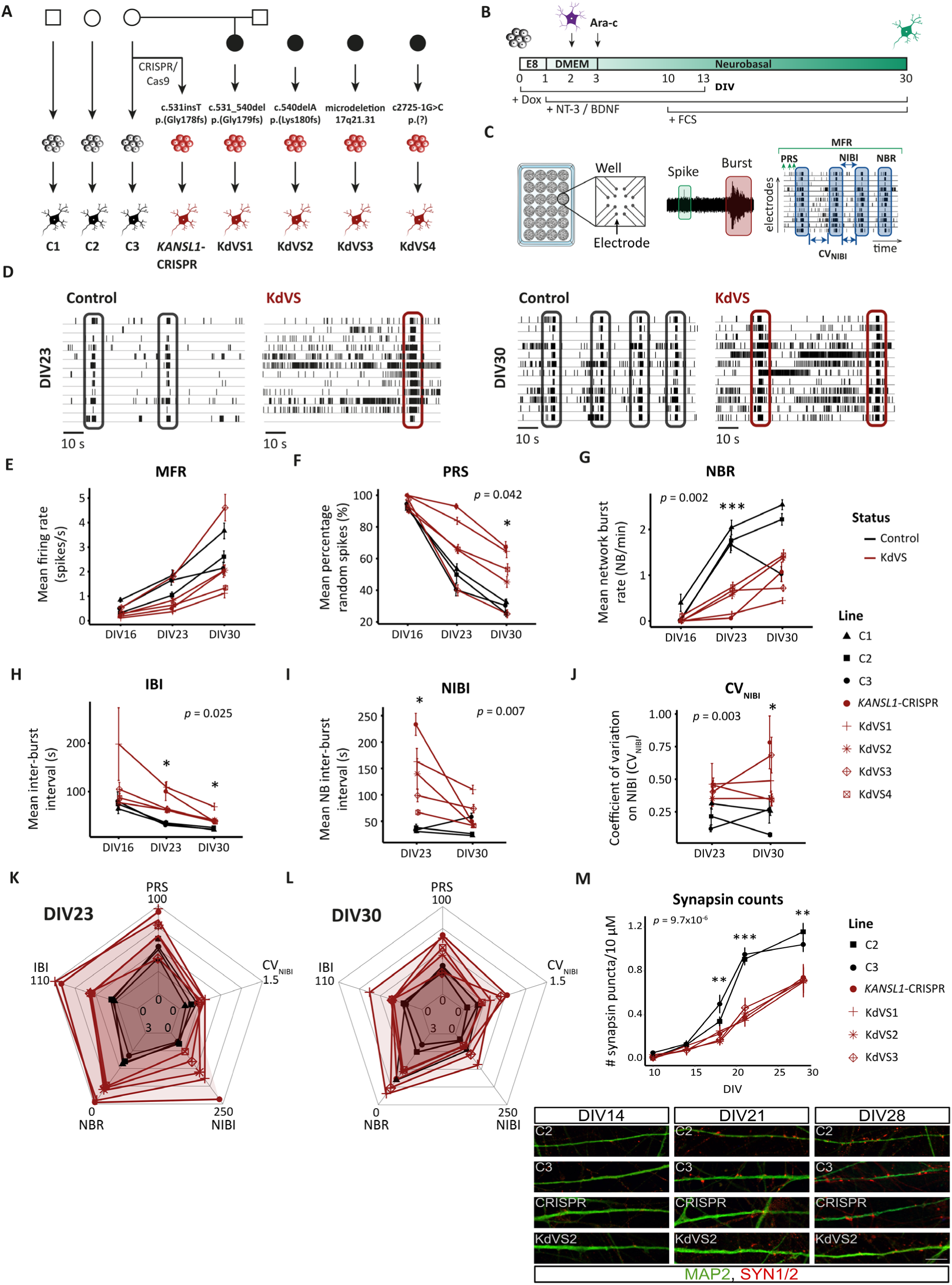
Impaired neuronal network function in hiPSC-neurons derived from KdVS individuals. **(A)** Overview of the cell lines generated from KdVS and controls. **(B)** Overview of neuronal differentiation protocol. Ngn2 expression is induced by adding doxycycline at DIV0. Rat astrocytes are added at DIV2 to support neuronal maturation, while neurotrophic factors (e.g. BDNF and NT-3) are added during differentiation. **(C)** Scheme of the microelectrode array (MEA) system, including a visual representation of parameters extracted from MEA measurements. **(D)** Representative raster plots showing neuronal network activity in a KdVS and a control line, at DIV23 and DIV30. Y-axis indicates the different electrodes, and X-axis indicates time. Each spike is represented by a vertical line, and network bursts are highlighted inside rectangles. **(E-J)** Plots showing mean value of MEA parameters describing neuronal network activity in KdVS (n_wells_=15-27) and control (n_wells_=15-22) hiPSC-derived neurons. Shapes represent different cell lines. Data is shown for DIV16, DIV23, and DIV30. **(K-L)** Radar charts showing the average value per line of PRS, IBI, NBR, NIBI and CV_NIBI_ in each condition at DIV23 **(K)** and DIV30 **(L)**. **(M)** Top: Number of synapsin puncta, observed per 10 µM of dendrite length, in KdVS (n_cells_=4-6) and control (n_cells_=4-6) hiPSC-derived neurons during differentiation. Bottom: Representative images of synapsin puncta staining in control and KdVS hiPSC-derived neurons. Significance was determined with one-way ANOVA on a linear model, followed by a t-test per individual time point. Data represent means ± SEM. \*\*\**p*-value <0.001, ** *p*-value <0.01, * *p*-value <0.05. CV_NIBI_ = coefficient of variation calculated on the NIBI; DIV = days *in vitro*; IBI = inter-burst interval; MFR = mean firing rate; NBR = network burst rate; NIBI = NB inter-burst interval; PRS = percentage of random spikes.

hiPSCs were plated on microelectrode arrays (MEAs) and differentiated into neurons by *Ngn2* overexpression for 30 days *in vitro* (DIV; **Fig. 1B**). The activity of the networks was measured during differentiation (**Fig. 1C**). Representative raster plots depicting the network phenotypes at DIV23 and DIV30 are shown in **Fig. 1D**. We calculated different “MEA parameters” to characterize the activity of the networks (**Fig. 1E-J**). In both control and KdVS neurons, the synchronized activity of the networks increased during differentiation, reflected by increased rate of single action potentials (mean firing rate, MFR), decrease in the percentage of random spikes (PRS), and higher network burst rate (NBR) over time (**Fig. 1E-G**). This indicated progressive maturation of the networks during differentiation. Nonetheless, differences in neuronal network activity features were observed for all KdVS lines compared to controls. The inter-burst interval (IBI; **Fig. 1H**), as well as the interval between network bursts (NIBI; **Fig. 1I**), were higher in KdVS neurons. Conversely, the NBR was reduced in KdVS (**Fig. 1G**). In addition, we found that the network burst activity in KdVS neurons was less regular, reflected by a higher coefficient of variation calculated on the NIBI (CV_NIBI_) (**Fig. 1J**).

Of note, we observed that the 17q21.31 microdeletion KdVS3 neurons, behaved differently in terms of MFR and PRS compared to the KdVS lines with a *KANSL1* mutation (KdVS1, KdVS2, KdVS4, *KANSL1*-CRISPR). KdVS lines with a *KANSL1* mutation significantly differed from controls on these parameters, with reduced MFR and increased PRS with respect to controls, while KdVS3 exhibited activity similar to control level (**Fig. 1E-F**)

In general, the largest differences in neuronal network activity were seen at DIV23 and to a lesser extent at DIV30, but differences remained present up to DIV79 (**Fig. 1K-L; Fig. S2A-C**), suggesting a general developmental delay in KdVS lines. Linear discriminant analysis (LDA) was used to classify the cell lines based on their neuronal network activity patterns (**Fig. S2D**). We observed high percentage of accurate predictions of the cell lines into their respective groups (control or KdVS) at both DIV23 and DIV30 (correct group classification DIV23: mean = 93%, DIV30: mean = 95%), whereas this was less clear at DIV16 (mean correct group classification = 67%), indicating that differences between KdVS and control lines occurred from DIV23 onwards (**Fig. S2E**).

To understand whether changes in synapse formation in KdVS neurons may contribute to the impairments in neuronal network activity, neurons were immunostained for synapsin1/2 over time. We found decreased synapsin1/2 puncta in all KdVS lines starting from DIV18, with largest differences observed at DIV21, consistent with the timing of more pronounced differences in neuronal network activity (**Fig. 1M**). Overall, despite some cell line-specific differences, common features of neuronal network dysfunction were identified in KdVS hiPSC-derived neurons, particularly for parameters describing the network burst activity.

### MEA-seq correlations identify increased CLC-4 contributing to reduced synchronized activity in KdVS neuronal networks

To investigate gene expression changes underlying the difference in neuronal network activity between KdVS and control neurons, we optimized a cost-efficient bulk RNA-seq method, which was then used in combination with MEA recordings in the same experiment (MEA-seq). We isolated RNA from neuronal cultures (**Fig. S3A**) directly after measuring the network activity at DIV30, from three controls (C1-3), the CRISPR line (*KANSL1*-CRISPR) and three KdVS lines (KdVS1-3). We also sequenced RNA samples from hiPSCs of the same lines (**Fig. S3B-C**). In both cell types, we observed decreased *KANSL1* expression activity (**Fig. S3D**). Surprisingly, we only identified 9 upregulated and 11 downregulated genes in KdVS neurons, as compared to controls (**Fig. S3E-F**; **Table S1)**. The presence of genotypic and phenotypic heterogeneity among KdVS individuals may hinder the identification of differentially expressed (DE) genes due to possible transcriptional heterogeneity within the KdVS cohort. More pronounced phenotypes are expected to be linked to more pronounced changes in gene expression. Since the transcriptional material was isolated in parallel from each recorded well of hiPSC-derived neurons, the MEA-seq approach brings the possibility to evaluate direct associations between gene expression and network activity changes. Thereby, MEA-seq allows to account for both genotypic and phenotypic heterogeneity between different lines. To integrate MEA and RNA-seq data, we calculated spearman correlations between the normalized number of transcripts of every gene and 10 MEA parameters describing the neuronal network phenotypes (**Table S2**). We used data from 20 samples that were included in both the MEA and RNA-seq experiments and that showed representative neuronal network phenotypes (C1-C3, KdVS1-3 and CRISPR-*KANSL1*; **Fig. S4A-F**). We excluded correlations caused solely by differences between KdVS and controls, or by gender, using a post-hoc analysis that only kept the correlations that remained significant within controls, KdVS, and female samples independently. The network burst duration (NBD) and the burst spike rate (BSR) correlated with the expression activity of a high number of genes (59-157 genes), whereas the PRS and the CV_NIBI_ correlated with only one gene (**Fig. 2A**). We identified 41 genes whose expression activity was significantly correlated with at least one of the affected activity parameters in KdVS (**Table S2**; **Fig. 2B**). Of these genes, 11 showed a trend for differential expression between KdVS and controls (*p*-value < 0.05; *C5orf49, CLCN4*, *DMD*, *RBBP7, TBX2, USP9X, FAM78B, TIMP2, TLE4, ZNF717* and *TIGD5*; **Fig. S4G**).

**Figure 2.**
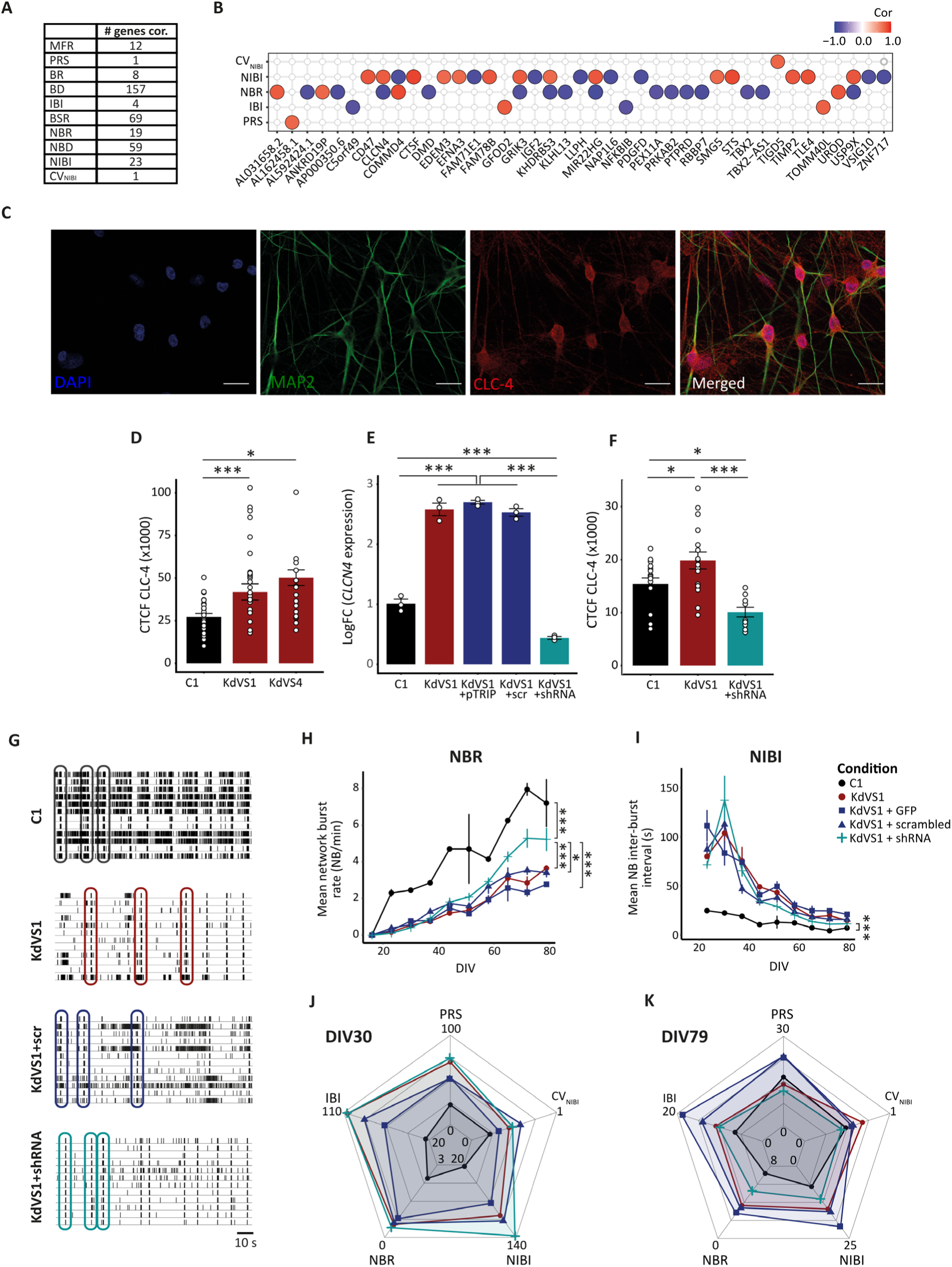
Increased *CLCN4* expression is linked to network dysfunction in KdVS hiPSC-derived neurons. **(A)** Table showing the number of significant correlations between the expression of individual genes and MEA parameters, after post-hoc analysis retaining the correlations that remained significant within controls, KdVS, and female samples independently. 20 wells of hiPSC-derived neurons at DIV30 were included in the MEA-seq experiment. **(B)** Correlation plot showing significant correlations (adj. *p*-value<0.05) between gene expression and 5 MEA parameters. **(C)** hiPSC-derived neurons (DIV30) were stained for DAPI (in blue), MAP2 (in green), and CLC-4 (in red). Scale = 25 µm. **(D)** Bar plot showing the CLC-4 corrected total cell fluorescence (CTCF) in C1 (n_cells_=24), KdVS1 (n_cells_=17) and KdVS4 (n_cells_=25) neurons. **(E)** Quantitative PCR (qPCR) of *CLCN4* in hiPSC-derived neurons (DIV30) from control (C1), non-treated KdVS1 and KdVS1 samples that followed transduction with an empty vector (KdVS1+pTRIP), a scrambled shRNA (KdVS1+scr), and a shRNA targeting *CLCN4* (KdVS1+shRNA). LogFC regarding C1 is shown for each condition, n=3 wells per condition. **(F)** Bar plots showing the CTFC for CLC-4 in C1 (n_cells_=17), KdVS1 (n_cells_=17) and KDVS1+shRNA (n_cells_=11) neurons. **(G)** Representative raster plots showing neuronal network activity in C1, KdVS1, KdVS1+scr, and KdVS1+shRNA hiPSC-derived neurons at DIV65. **(H-I)** Linear plots showing the NBR **(H)** and the NIBI **(I)** in C1 (n_wells_=2), KdVS1 (n_wells_=3), KdVS1+pTRIP.GFP (n_wells_=4), KdVS1+scr (n_wells_=4), and KdVS1+shRNA (n_wells_=5). Data represent mean ± SEM and are shown for DIV16-79. Shapes represent different cell lines or conditions. Significance was determined by one-way ANOVA, followed by a Tukey HSD post-hoc test. Significant effects are only shown for comparisons between all conditions and the KdVS1+shRNA (\*\*\**p*-value<0.001, ** *p*-value<0.01, * *p*-value <0.05). **(J-K)** Radar charts showing the average PRS, IBI, NBR, NIBI and CV_NIBI_ in C1, KdVS1, KdVS1+pTRIP.GFP, KdVS+scr, and KdVS+shRNA neurons at DIV30 **(J)** and DIV79 **(K)**. BD = burst duration; BR = burst rate; BSR = burst spike rate; CV_NIBI_ = coefficient of variation calculated on the NIBI; DIV = days *in vitro*; IBI = inter-burst interval; MFR = mean firing rate; NBD = network burst duration; NBR = network burst rate; NIBI = NB inter-burst interval; PRS = percentage of random spikes; logFC = log fold change.

The expression of *CLCN4*, a gene encoding the voltage-dependent chloride channel CLC-4, was negatively correlated with the NBR (cor = -0.85, adj. *p-*value = 0.02) and positively correlated with the NIBI (cor = 0.80, adj. *p-*value = 0.01; **Fig. 2B, Fig. S4H-I**). Immunocytochemistry staining of CLC-4 localized it in the soma and dendrites of hiPSC-derived neurons (**Fig. 2C**), and confirmed increased expression of CLC-4 in KdVS neurons (**Fig. 2D**). CLC-4 showed increased expression in KdVS4 neurons as well, which were not included in the MEA-gene correlation analysis but exhibited decreased NBR and increased NIBI compared to controls too (**Fig. 2D**). Independent qPCR analysis of *CLCN4* also confirmed increased expression activity in KdVS neurons (**Fig. 2E**). To test whether the correlation between *CLCN4* expression and NBR represents a causal relationship, we performed a knockdown experiment using a *CLCN4* shRNA in KdVS1 hiPSC-derived neurons, and investigated the effect of reducing *CLCN4* expression on the neuronal network phenotype. We confirmed that *CLCN4* shRNA treatment decreased both *CLCN4* gene and CLC-4 protein expression towards control level (**Fig. 2E-F; Fig. S4J**). In contrast, neither the empty vector nor the scrambled shRNA (control-treatment) affected *CLCN4* expression (**Fig. 2E**). *CLCN4* knockdown in KdVS neurons increased the NBR over time as compared to control-treated KdVS neurons (**Fig. 2G-H**). The strongest effect was observed from DIV65 onwards. This result confirms a causal relationship between increased *CLCN4* expression and reduced NBR in KdVS hiPSC-derived neurons. The effect of *CLCN4* knockdown on NIBI did not differ from control-treatment until DIV65 (**Fig. 2I**). Starting from DIV65, the NIBI in neurons treated with *CLCN4* shRNA was significantly lower compared to control-treatment (untreated KdVS1: adj. *p* = 0.01; empty vector KdVS1: adj. *p* < 0.001; scrambled shRNA KdVS1: adj. *p* = 0.007), and thus, more similar to the NIBI in control networks. Of note, the other activity parameters were not significantly affected following *CLCN4* shRNA treatment (**Fig. 2J-K**). Overall, these results suggest that altered expression of *CLCN4* causally contributes to the abnormal neuronal activity in KdVS neurons, in terms of NBR and NIBI. It is worth noting that downregulation of *CLCN4* expression had an impact on the network activity during later stages of differentiation and did not completely restore the network phenotype, suggesting that additional mechanisms contribute to the network bursting phenotype in KdVS.

### Impaired mitochondrial function is linked to the neuronal network phenotype of KdVS

To explore a potential contribution of other biological processes to alterations in neural network activity, we performed GSEA on genes ranked according to their correlation with each of the 10 MEA parameters (adj. *p*-value < 0.05; **Table S2**). These gene sets encompassed terms such as “nonsense-mediated decay”, “SLIT-ROBO signaling”, “aggrephagy”, “ciliary function”, and protein “translation”, “localization” and “ubiquitination” (**Fig. S4K**).

We identified enriched gene sets associated with the MEA parameters affected in KdVS. For instance, “axon development”, “synapse organization”, “GTPase signaling”, and “neuron projection development” were significantly associated with parameters that describe bursting activity, such as BR, IBI, NBR, and NIBI (**Fig. 3A**). Furthermore, several HPO terms were also enriched for the MEA parameters affected in KdVS, including terms associated with epileptic seizures, intellectual disability, and muscle hypotonia (**Fig. 3B**), which are well-known clinical hallmarks of KdVS

**Figure 3.**
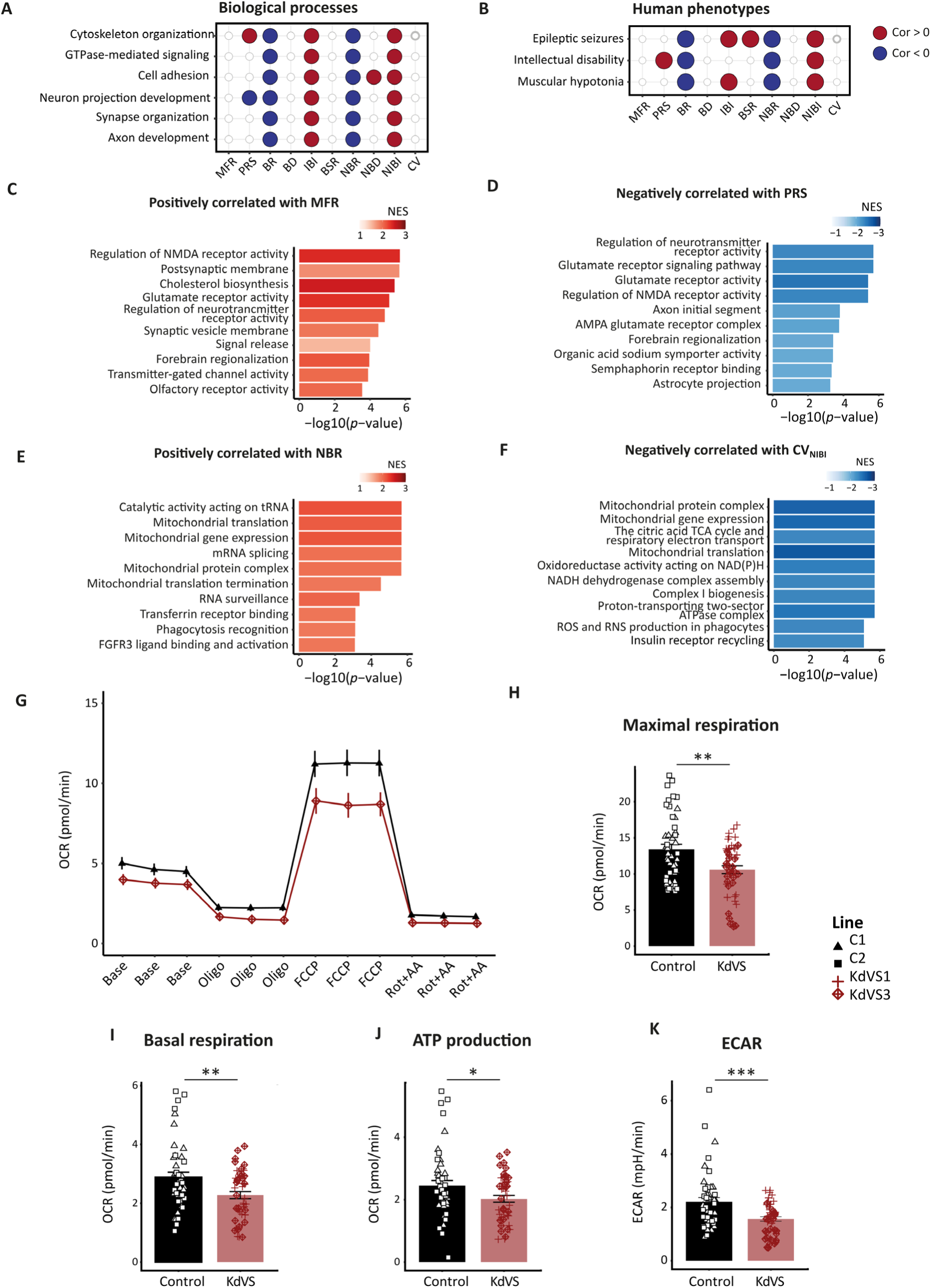
Changes in the expression of synaptic-related and mitochondrial genes are linked to changes in the activity of hiPSC-derived networks. **(A-B)** Correlation plot showing gene sets significantly enriched (adj. *p*-value<0.05) for genes that either positively or negatively correlated with the MEA parameters, for Gene Ontology (GO) biological processes **(A)** and Human Phenotype Ontology (HPO) terms **(B)**. **(C-D)** Bar plots showing top Reactome pathways and GO terms enriched for genes positively correlated with the MFR **(C)**, and for genes negatively correlated with the PRS **(D)**. **(E-F)** Bar plots showing top Reactome pathways and GO terms enriched for genes positively correlated with the NBR **(E)** and for genes negatively correlated with CV_NIBI_ **(F)**. Data was generated from control (n_wells_=9) and KdVS (n_wells_=11) hiPCs-neurons at DIV 30. **(G-K)** Assessment of mitochondrial function. **(G)** OCR measurements at basal level (base), following treatment with 2 µM oligomycin (oligo), 3 or 4 µM FCCP, and 0.5 µM rotenone and antimycin A (Rot+AA), performed in hiPSC-derived neurons from control (C1, C2; n_wells_=23-24) and KdVS lines (KdVS1, KdVS3; n_wells_=24) at DIV7. **(H-K)** Mitochondrial function by means of maximal respiration **(H)**, basal respiration **(I)**, ATP-linked respiration **(J)**, and ECAR **(K)**, which represents the glycolysis rate. The ECAR was calculated by averaging the ECAR values over the first three basal measurements. Shapes indicate the cell line. All measurements were normalized to CS activity. Significance was determined by a one-sample *t*-test. Data represent means ± SEM. *** *p*-value <0.001, ** *p*-value <0.01, * *p*-value <0.05. CV_NIBI_ = coefficient of variation calculated on the NIBI; MFR = mean firing rate; NBR = network burst rate; NES = normalized enrichment score; NIBI = NB inter-burst interval; PRS = percentage of random spikes, ECAR = extracellular acidification rate; OCR = oxygen consumption rate; CS = citrate synthase.

Interestingly, we identified biological processes linked to MEA parameters affected in KdVS that were not captured with GSEA on the DE genes. For instance, the gene sets ranked highest in enrichment for genes exhibiting a positive correlation with the MFR and negative correlation with the PRS were terms associated with glutamatergic receptor activity and synaptic signaling-related genes (**Fig. 3C-D**). On the other hand, genes that displayed a positive correlation with NBR were enriched in gene sets associated with mitochondrial gene expression and translation (**Fig. 3E**). These gene sets were negatively related to CV_NIBI_ (**Fig. 3F**). These results suggest that mitochondrial function is important for synchronous network activity, which is decreased in KdVS neurons *in vitro* and may be impaired in KdVS. We therefore next evaluated mitochondrial function in hiPSC-derived neurons from KdVS and controls with a Seahorse XF Cell Mito stress test. We measured the oxygen consumption rate (OCR), which measures mitochondrial respiration, and the extracellular acidification rate (ECAR), which reflects glycolytic functioning. To determine the rates per mitochondrion, both parameters were normalized to citrate synthase (CS) levels [36]. We observed decreased basal and maximal respiratory capacity (induced by FCCP treatment) in KdVS neurons compared to controls (**Fig. 3G-I).** The addition of oligomycin, an ATP synthase inhibitor, demonstrated that KdVS neurons were also not efficient as control neurons in ATP production (**Fig. 3J**). Lastly, the ECAR levels were decreased in KdVS neurons, which is indicative of decreased glycolytic activity (**Fig. 3K**). Overall, these data demonstrate that mitochondrial function and metabolic activity are decreassed in KdVS neurons *in vitro*. This likely contributes to impaired neuronal network function in KdVS.

### Computational drug screening based on transcriptional changes in KdVS neurons

After identifying the transcriptomic signature and probable molecular mechanisms responsible for impaired neuronal network functioning in KdVS, our next step was to search for compounds that could potentially reverse these molecular changes and consequently restore neuronal activity patterns. To this end, we investigated compounds that could counteract the transcriptional changes associated with KdVS neurons (DIV30), based on transcriptomic signatures from the LINCS database. Considering the differences in the transcriptional profile of different KdVS cell lines, the query genes were selected by identifying DE genes between KdVS and corresponding control lines, such as *KANSL1*-CRISPR and its isogenic control (C3), and between KdVS1 and its parental control (C3), identifying 201 and 1060 significant DE genes, respectively (adj. *p*-value < 0.05; **Fig. 4A-B**; **Table S3**; Supplementary methods). Gene expression profiles of KdVS2 and KdVS3 were compared to pooled C1 and C2, and resulted in the identification of 221 and 376 significant DE genes, respectively (**Fig. 4B**; **Table S3**). For each cell line, the top 150 upregulated and top 150 downregulated were selected to query the LINCS database (**Fig. 4A-B**). There was little overlap between the query genes of different lines, with 45-85% unique query genes per line (**Fig. 4C**). The greatest overlap was observed between KdVS1 and the *KANSL1*-CRISPR, which share ∼50% of their genetic background.

**Figure 4.**
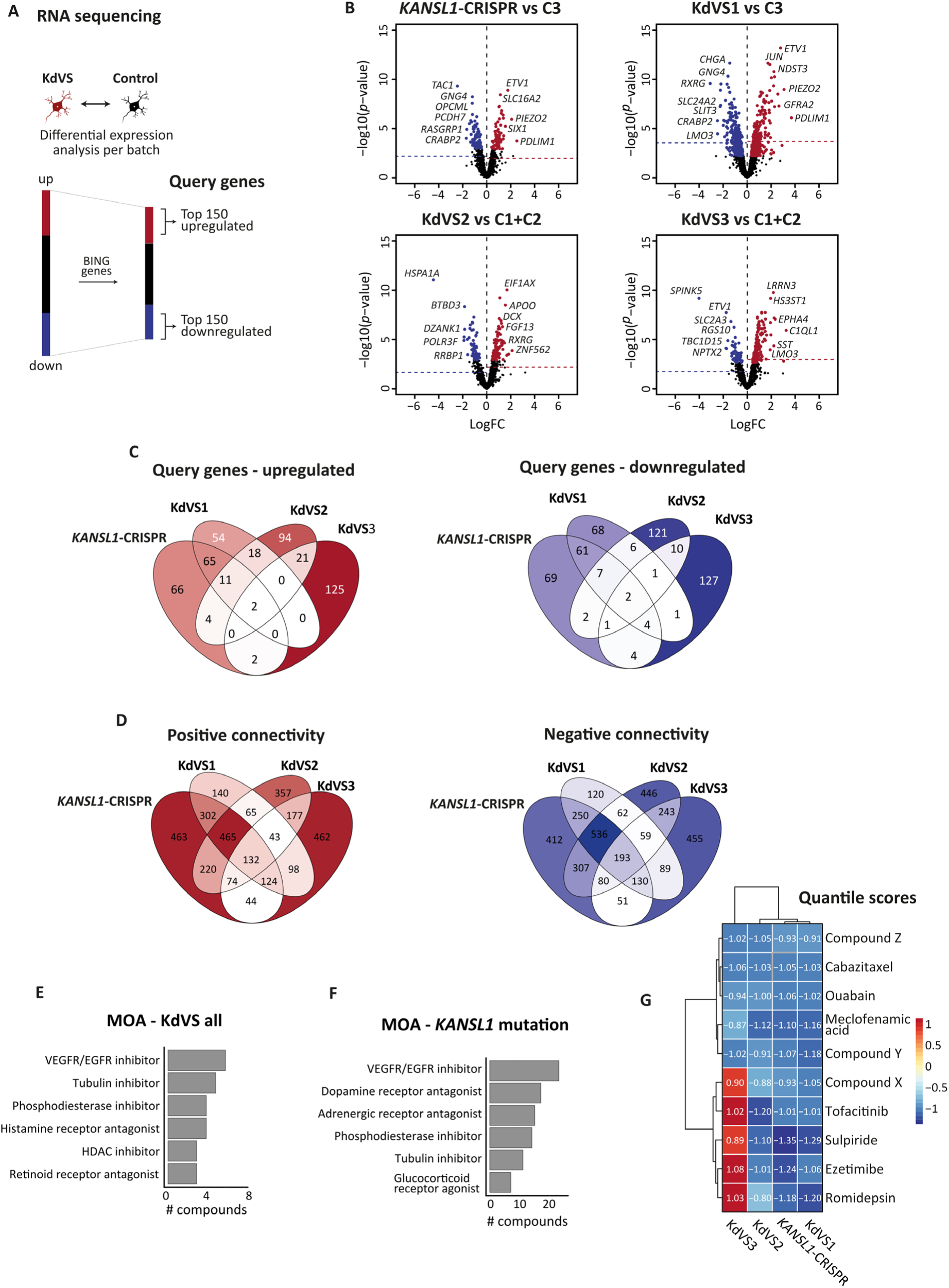
Computational drug screening based on the transcriptional signature of KdVS neurons. **(A)** Schematic showing the steps to define query genes. Differential expression (DE) analysis between KdVS and control neurons (DIV30) was performed per batch. Subsequently, DE genes were filtered for the BING (Best Inferred Genes) of the LINCS database. **(B)** Volcano plots showing DE genes in *KANSL1*-CRISPR vs C3 (top left), KdVS1 vs C3 (top right), KdVS2 vs C1+C2 (bottom left) and KdVS3 vs C1+C2 (bottom right). Genes with adj. *p*-value<0.05 are labelled. The horizontal dashed line represent the selection threshold for query genes per line (top 150 down- and upregulated genes). **(C)** Overlap of query genes between the different KdVS cell lines. **(D)** Overlap of drug perturbations with positive (left) and negative (right) connectivity between individual KdVS queries. **(E-F)** Barplot depicting the top shared mechanisms of action (MOA) of compounds that showed negative connectivity with all KdVS lines (n_total_ = 193) **(E),** and with KdVS lines with a *KANSL1* mutation (*KANSL1*-CRISPR, KdVS1, and KdVS2; n_total_ = 729) **(F)**. **(G)** Heatmap showing quantile scores for the top 10 compounds per KdVS line, with rows and columns ordered based on hierarchical clustering. LogFC = log-fold change.

Out of 30,780 perturbations characterized in the LINCS database, we identified ∼2,500-4,500 as potentially suitable for modifying the expression of the query genes, across the different KdVS cell lines (**Fig. S5A**). Compounds that were predicted to alter the expression of query genes in the same direction as the disease effect were classified as having positive connectivity. Conversely, compounds exhibiting negative connectivity were those predicted to modulate gene expression in the opposite direction to the disease effect. Therefore, we had particular interest in compounds with a negative connectivity, since these may reverse the KdVS transcriptomic signature. Despite the little overlap between query genes (**Fig. 4C**), there was high overlap in compounds with potential therapeutic effect across the different KdVS lines (**Fig. 4D**). This finding suggests that similar mechanisms may be disrupted across the different cell lines. The overlap was higher between KdVS lines with a *KANSL1* mutation (*KANSL1*-CRISPR, KdVS1, and KdVS2). Out of 2,305 overlapping compounds, 132 showed positive connectivity, while 193 compounds showed negative connectivity with all four KdVS signatures (**Fig. 4D**). We selected compounds for experimental validation within the ones presenting negative connectivity with at least all *KANSL1* mutation lines. We used two further selection criteria; 1) compounds representing a class of compounds with shared mechanisms of action (MOA) and 2) compounds with the most negative quantile scores. In addition, we only considered compounds already launched to clinical phase. The top common MOA included VEGFR/EGFR inhibitors, tubulin inhibitors, phosphodiesterase inhibitors, histamine receptor antagonists, HDAC inhibitors, and retinoid receptor antagonists (**Fig. 4E-F**). Dopamine and adrenergic receptor antagonists, and glucocorticoid receptor agonists were identified specifically for the *KANSL1* mutation lines (**Fig. 4F**). Based on the top MOAs, we selected cabazitaxel (microtubule inhibitor), romidepsin (HDAC inhibitor), and prednisone (glucocorticoid receptor agonist). Based on the top negative connectivity score (Q KdVS all< -0.9 or Q KANSL1 < -1.0), we selected ezetimibe (cholesterol inhibitor), meclofenamic acid (cyclooxygenase inhibitor), ouabain (ATPase inhibitor), sulpiride (dopamine receptor antagonist), tofacitinib (JAK inhibitor), phloretin (sodium/glucose co-transporter inhibitor), and fasudil (Rho-associated kinase inhibitor). Details about the compounds can be found in **Table S4**. Ezetimibe, romidepsin, sulipride, tofacitinib, and prednisone showed negative connectivity with the *KANSL1* mutation lines, but not with KdVS3, while the other compounds were negatively connected with all KdVS cell lines (**Fig. 4G**).

### Screening the effect of selected compounds on neuronal network activity identified three potential candidates for KdVS treatment

We first tested the effect of the selected compounds (**Table S4**) on neuronal network activity in *KANSL1*-CRISPR neurons. We treated hiPSC-derived neurons from DIV6 onward (**Fig. S6A**), with the same concentration used for the phase I LINCS L1000 study (10 µM). Since the compounds were selected based on transcriptomic changes at DIV30, we focused on their effect on network activity at DIV30. Cabazitaxel and romidepsin treatment induced cell death (**Fig. S6B**). Ouabain treatment also had a negative effect on the neurons, reflected in reduced cell density and null network activity (**Fig. S6C**). Lowering their concentration to 100 nM was not sufficient. They remained toxic, and were excluded from further analysis. Tofacitinib treatment (10 µM) induced some cell death (**Fig. S6D**), but the remaining neurons were electrically active. The other five compounds did not affect cell density and were not toxic (**Fig. S6D**). We then investigated whether these compounds were able to reverse the neuronal network phenotype of KdVS to control level, by increasing MFR and NBR, and decreasing the PRS, IBI, NIBI, and CV_NIBI_.

At DIV30, and based on individual activity parameters, none of the compounds fully reverted the activity phenotype of *KANSL1*-CRISPR networks to control levels (**Fig. S6E-J**). Meclofenamic acid, ezetimibe, tofacitinib, and sulpiride exacerbated the disease effect, showing a tendency to induce a more desynchronized network and increase the IBI (**Fig. S6E-F**). Conversely, prednisone demonstrated more promising effects, with a trend toward improving overall spiking activity by increasing the MFR (**Fig. S6G**) and significantly increasing the NBR (**Fig. S6H**), along with a trend toward decreasing the NIBI (**Fig. S6I**). Phloretin also showed a trend toward increasing the NBR and reducing the NIBI and CV_NIBI_. Fasudil exhibited a tendency to improve the regularity of network bursting activity by reducing the CV_NIBI_ (**Fig. S6J**). Given that prednisone, phloretin, and fasudil showed the most positive results, we tested their effects on a independent cell line. We employed CRISPR/Cas9 on C1 to introduce a nonsense mutation in exon 2 of *KANSL1*, generating *KANSL1*-CRISPR_C1. *KANSL1*-CRISPR_C1 segregated from its control in a similar pattern to *KANSL1*-CRISPR, as shown by LDA on PRS, IBI, NBR, and CV_NIBI_ (**Fig. 5A**). C1 control networks exhibited synchronised NBs at DIV28, while *KANSL1*-CRISPR_C1 networks showed a more desynchronised network with longer and more complex NBs (**Fig. 5B**). For this round of experiments, we took along both the mutated line and its isogenic control, allowing a direct comparison between the KdVS-compound-treated and control networks. We tested two different doses (0.3 µM and 1 µM) and employed a multiparametric approach, including UMAP and clustering analysis, to explore their effects along the developmental trajectory. After 24 hours of drug exposure, the KdVS segregated from the control line in the UMAP space, showing no general effects of the drugs on KdVS-related activity (**Fig. S7A**). Hierarchical clustering identified two activity clusters: one associated with control activity and the other with KdVS (**Fig. S7B**). We evaluated the association between conditions (control, KdVS, and compound-treated KdVS) and each activity cluster using a Bayesian generalized linear model, which estimates the odds of a network being assigned to each cluster and provides 90% credible intervals for the estimates. Notably, after approximately 2 weeks of treatment (12 to 14 days), the treated networks shifted towards the control activity cluster (**Fig. 5C-E**), indicating that a longer time period was needed for the drugs to exert their effects, likely due to the transcriptional nature of the changes. The strongest effect was observed with phloretin at DIV28 (**Fig. 5C**), when both concentrations induced a significant change for the KdVS-treated networks towards the control phenotype, with 80% of phloretin-treated KdVS networks assigned to the control activity phenotype. In particular, the IBI was significantly increased for both concentrations of phloretin, more closely representing the IBI for C1, while the IBI-CoV was significantly decreased only for the higher concentration. A similar effect was observed with the higher concentration of fasudil (1 µM) at DIV28 (**Fig. 5D**). Here, both the IBI and the IBI-CoV were significantly increased only at the higher concentration of fasudil. The most pronounced effect of prednisone was observed at a later time point (DIV30; **Fig. 5E**). Although increasing the concentration resulted in a higher percentage of KdVS-treated networks being assigned to the control cluster (**Fig. 5E.II**), there was no significant shift towards the control cluster for prednisone-treated KdVS networks (**Fig. 5E.III**). This suggests that a higher concentration of prednisone may be necessary to effectively rescue KdVs activity.

**Figure 5.**
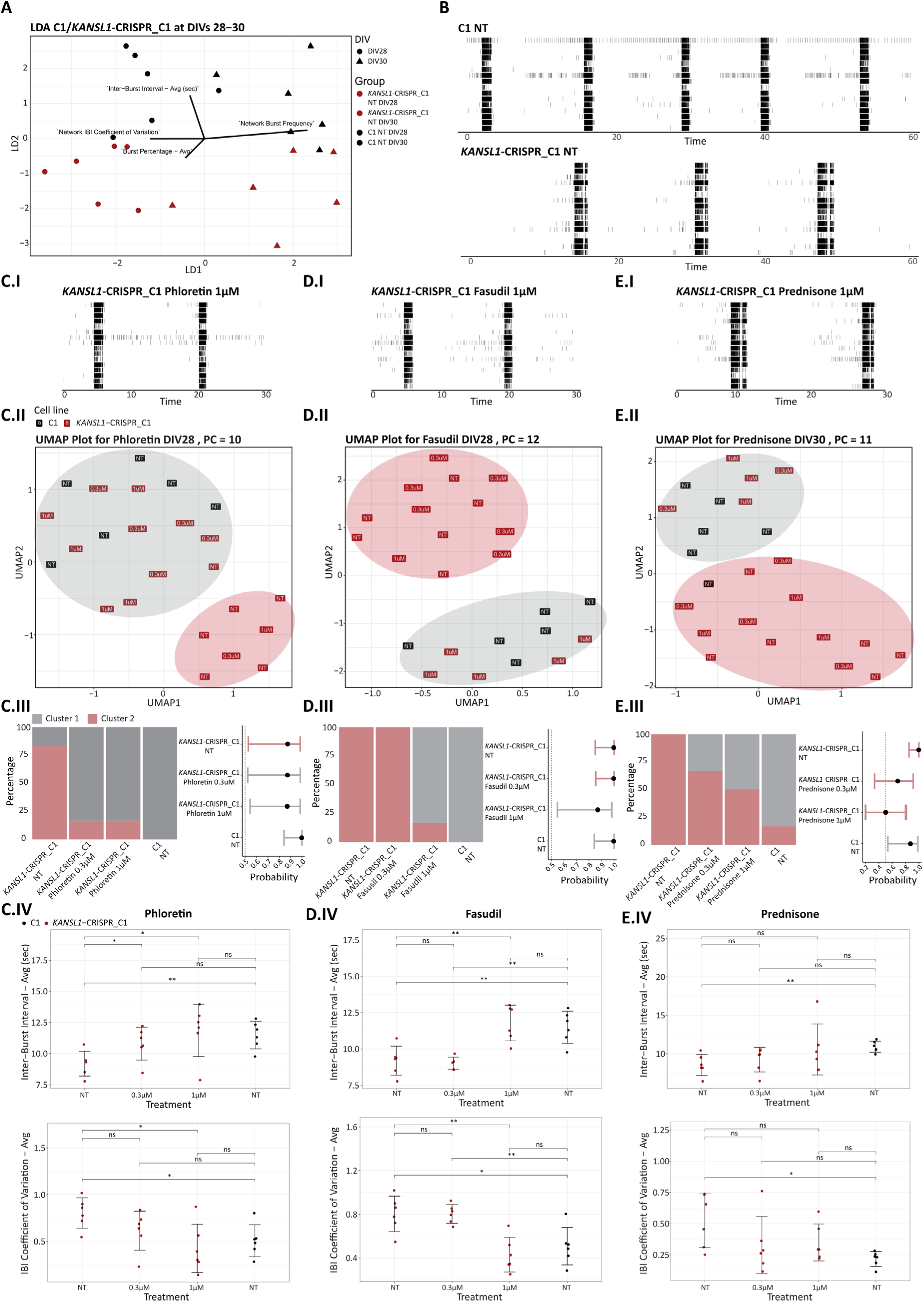
Effect of fasudil, phloretin and prednisone on neuronal network activity in *KANSL1*-CRISPR_C1-derived neurons. **(A)** Linear discriminant analysis (LDA) performed on 4 MEA parameters affected in KdVS (PRS, IBI, NBR, and CV_NIBI_), for *KANSL1*-CRISPR_C1 (red) and its isogenic control C1 (black) at DIVs 28 and 30. The arrows indicate the direction of MEA parameters in KdVS. Shapes represent different DIVs. **(B)** Representative rasterplots for control C1 and *KANSL1*-CRISPR_C1 at DIV28 with no treatment. **(C-E)** Compounds were added at final concentrations of 0.3 µM and 1 µM. Representative rasterplots are shown for 1µM treatment of phloretin **(C.I)**, fasudil **(D.I)** and prednisone **(E.I)**. UMAPs were performed on principal components (PCs) obtained from network activity parameters for C1 and *KANSL1*-CRISPR_C1-derived neurons treated with phloretin (**C.II**; DIV28; 10 PCs), fasudil (**D.II**; DIV28; 12 PCs), and prednisone (**E.II**; DIV30; 11 PCs). Subsequently, we performed hierarchical clustering analysis based on the two UMAP dimensions. The ellipses group the recordings assigned to each cluster. Proportion of recordings assigned to each hierarchical clustering per condition are depicted for phloretin in **(C.III)**, fasudil **(D.III)** and prednisone **(E.III)**. The association between condition and activity cluster was tested using a Bayesian generalized linear model with a binomial distribution and a logit link function. The probability plots in **(C.III-E.III)** show the 90% credible intervals for the odds of a network being assigned to each cluster under each condition, converted into probabilities. **(C.IV-E.IV)** Inter-burst-interval (top) and inter-burst-interval (IBI) coefficient of variation (bottom) at DIV28 for phloretin, fasudil and prednisone. The difference between non-treated and treated conditions was tested using an unpaired *t*-test; * *p*-value< 0.05, ** *p*-value < 0.01.

In summary, fasudil and phloretin emerged as the most promising candidates, consistently improving KdVs-related activity across different KdVS cell lines.

## Discussion

For the present study, our aim was to conduct an integrated analysis of molecular and electrophysiological changes in KdVS neurons. We designed a novel approach that combined transcriptome profiling with network activity measurements (MEA-seq), to identify potential molecular changes underlying the variation in network activity associated with KdVS. Through MEA-seq and knockdown experiments conducted on KdVS and control hiPSC-derived neurons, we demonstrated *CLCN4* as candidate gene modulating the aberrant network activity phenotype exhibited by KdVS neurons characterized by reduced synchronized bursting activity. Additionally, using MEA-seq we identified impaired mitochondrial function likely related to the network phenotype of KdVS neurons. This was subsequently confirmed by experimental assessment of mitochondrial function. In pursuit of druggable pathways counteracting the relevant transcriptional changes and thus restoring the neuronal activity patterns in KdVS, we computationally screened drug perturbation signatures in the LINCS Consortium database, using the transcriptomic profile of KdVS derived neurons as a query. From more than 700 compounds predicted to reverse the transcriptomic profile of KdVS neurons, we selected 10 for experimental validation, ultimately identifying two candidates with potential for ameliorating the impaired network activity in KdVS. This work introduces a new approach to study neurodevelopmental disorders and may strengthen the development of effective drug treatments, by dissecting the molecular mechanisms underlying neural network dysfunction.

Knockdown of *CLCN4* in KdVS neurons *in vitro* partially restored the synchronized bursting activity to control level without modifying other activity parameters. This highlights the ability of the MEA-seq to identify associations between individual genes and specific activity parameters, providing the opportunity to fine-tune particular activity features without necessarily engaging all forms of network activity. *CLCN4* encodes for the chloride voltage-gated channel 4 (CLC-4), which is part of the CLCN family consisting of 9 members. Interestingly, mutations in *CLCN4* leading to loss of function of CLC-4 have been linked to epilepsy [41] and intellectual disability syndrome [42]. CLC-4 localizes at the endoplasmic reticulum [43] and the endosomes [44], and is highly expressed in the brain, heart and muscle tissue [45]. We have previously shown a higher presence of autophagosomes and decreased lysosome function in KdVS neurons *in vitro* [6], which combined with increased CLC-4 expression may indicate impaired function of the endolysosomal system in KdVS. Autophagy and the endolysosomal system are both involved in neurodevelopment, by regulating neuronal protein turnover, particularly at the presynapse [46]. The endolysosomal system and mitochondrial function are interconnected as well, and impaired function of one system influences the functioning of the other [47].

We identified a positive correlation between mitochondrial gene expression and parameters describing network burst activity. Mitochondrial ATP production is important for synaptic functioning [48, 49], and mitochondrial dysfunction has been reported in a large number of neurodevelopmental and neurodegenerative diseases, including Down syndrome [50], Parkinson’s disease [51], Alzheimer’s disease [52], schizophrenia, bipolar disorder, as well as autism spectrum disorder [53, 54]. In addition, impaired mitochondrial function has been specifically linked to neuronal dysfunction in a *in vitro* model of Huntington’s disease [55], epilepsy [56], and mitochondrial encephalomyopathy, lactic acidosis, and stroke-like episodes (MELAS) [23]. We corroborated an impaired mitochondrial function in KdVS neurons *in vitro*, demonstrated by reduced basal and maximal respiration, suggesting that mitochondrial dysfunction may be contributing to altered neuronal function and neurodevelopment in KdVS patients. Additionally, the tissues affected in KdVS are all high energy-dependent, including the brain, heart and muscle tissue [1], further suggesting an involvement of mitochondrial dysfunction underlying KdVS pathophysiology. Future studies tuning mitochondrial activity, autophagy, and the endolysosomal system may help to understand how these processes contribute to the neuronal network phenotype of KdVS neurons.

The comparative analysis on the RNA-seq data including all KdVS lines and controls identified low numbers of differentially expressed genes, probably due to the heterogeneity across the different KdVS cell lines, and neither *CLCN4* nor mitochondrial gene expression changes were detected. A potential driver of heterogeneity may be the difference in haplotype between the cell lines. Duplications can occur in the 17q21.31 region, giving rise to different *KANSL1* fusion transcripts depending on the H1 or H2 haplotype [57]. These fusion transcripts only encode for the coiled-coil domain of KANSL1 but lack the PEHE domain, therefore may interfere with the function of KANSL1. Presence or absence of such fusion genes, in combination with loss of function of KANSL1, could possibly explain the variability in gene expression profiles of the different KdVS lines. The advantage of MEA-seq is that no binary group labels are used to differentiate between the samples. Instead, a continuous scale describing the functional phenotype is considered to identify the changes in gene expression activity, thereby taking into account the phenotypic heterogeneity of the cell lines. This might be especially relevant for disorders involving high genotypic and phenotypic heterogeneity between cases, such as KdVS [26]. For instance, we observed differences in the neuronal network phenotype of the KdVS cell line with a 17q21.31 microdeletion, as compared to the cell lines with a mutation in *KANSL1*. While the cell lines with a mutation in *KANSL1* presented reduced mean firing rate (MFR) and higher percentage of random spikes (PRS), both parameters describing spiking activity, the cell line harbouring a microdeletion behaved similar to controls in terms of these parameters. MEA-seq identified decreased expression of genes related to synaptic signaling linked to reduced MFR and higher PRS, suggesting that synaptic signaling may only be affected in KdVS neurons with a *KANSL1* mutation. However, we only included one cell line with a 17q21.31 microdeletion and more donors may be included to corroborate a genotype-phenotype relationship.

To experimentally assess the transcriptome’s ability to predict network functionality and with the aim to restore KdVS-related activity, we performed a computational screening to identify compounds that could correct the transcriptomic signature and, consequently, the network activity in KdVS. We selected compounds for subsequent experimental assay based on their predicted ability to counteract the gene expression signature across the majority of KdVS lines. This approach aims to target the underlying mechanisms of KdVS pathology and may offer several advantages over symptomatic treatment. These compounds may have the potential to prevent the onset of symptoms rather than alleviating existing symptoms, and could have a positive long-term impact on the disease course. Of the ten compounds tested, phloretin, a sodium/glucose co-transporter inhibitor, and fasudil, a Rho-associated kinase inhibitor, consistently improved the activity in KdVS networks towards the control phenotype. Rho kinase (ROCK) inhibitors have a wide variety of therapeutical applications, including applications in neurodegenerative disorders [58]. KdVS-derived neurons exhibited increased expression of genes related to Rho GTPase signaling, therefore, ROCK inhibition likely compensated for this overactivation. In addition, ROCK inhibition was shown to increase synapse formation and to regulate autophagy in neurodegenerative disorders [59]. Since synapse formation and autophagy seem to be altered in KdVS, these could be the mechanisms by which fasudil improved the activity of the networks. On the other hand, sodium/glucose co-transporter inhibitors mainly target SGLT1 and SGLT2, and are often used as therapy for diabetes [60]. However, these transporters are not expressed in the brain, therefore phloretin likely improved the KdVS neuronal network phenotype through other targets. Additional experiments should be conducted to explore the mechanisms of action underlying the functional improvements facilitated by the compounds. For example, RNA-seq analysis of the compound-treated samples could offer insights into the potential molecular changes reversed by the compounds, whereas quantifying the number of synapses might help elucidate relevant cellular-level alterations associated with network functioning.

A potential limitation of this study is that the compounds were tested on neurons co-cultured with rat astrocytes, while the selection of compounds was based on their predicted ability to improve gene expression changes specifically in neurons. We cannot exclude the possibility that an indirect effect of the compounds on the rat astrocytes contributed to the changes in network activity, since neuronal function is highly dependent on astrocytes [61]. Performing RNA-seq experiments on the co-cultures following drug exposure would provide more insight into the cell type-specific contributions. Additionally, a more complex model system, such as brain organoids, including astrocytes derived from patients, may offer a more suitable setup for testing cell type-specific effects. Another limitation is the relatively low number of biological and technical replicates included in the drug assay. Follow-up experiments with larger sample sizes, encompassing multiple patient lines and dose-response curves, are still necessary to assess the effects in a patient-specific background and at clinically-relevant concentrations.

Summarizing, the MEA-seq analysis provided insights into the underlying mechanisms of KdVS by accounting for phenotypic heterogeneity between the samples. Through a computational approach and a pre-screening on MEAs, we successfully identified compounds that improved the network activity of KdVS neurons *in vitro*. The MEA-seq approach combined with the drug identification pipeline holds potential for unraveling the mechanisms responsible for neuronal network dysfunction and identifying candidate compounds for the treatment of different neurodevelopmental disorders. Optimization of the MEA-seq technology, including a variety of relevant neuronal and glial cells would significantly advance our comprehension of the mechanisms governing specific aspects of neuronal network activity in both normal and altered neurodevelopment.

## Acknowledgments

This work was supported by a grant from the Radboud university medical center and Donders Institute for Brain, Cognition and Behaviour (B.d.V.) and SFARI grant 890042 (to N.N.K.). This work was partially funded from the European Union’s Horizon 2020 research and innovation programme under grant agreement N°825575 (The European Joint Programme Rare Diseases, EJP RD) and from the Dutch Research Council (NWO) through a grant to the Netherlands X-omics initiative (project 184.034.019).

## Data availability

All data and codes generated in this study are available from the corresponding author upon request. This study did not generate any novel reagents apart from hiPSC lines. The cell lines used in this study are available upon request with a completed Materials Transfer Agreement from the corresponding author.

