## Supplementary information and figures for "Integrative transcriptomics and electrophysiological profiling of hiPSC-derived neurons identifies novel druggable pathways in Koolen-de Vries Syndrome"

Supplementary methods:

*hiPSCs generation*

C1 and C2 control cell lines were reprogrammed with episomal vector-based expression of the transcription factors: *Oct4*, *c-Myc*, *Sox2*, and *Klf4* [1]. Fibroblasts of C3 and KdVS1 were reprogrammed using the Simplicon™ reprogramming kit (Millipore), with *Oct4*, *Klf4*, *Sox2*, and *Glis1*. KdVS2 and KdVS4 fibroblasts were reprogrammed by lentiviral-mediated overexpression of *Oct4*, *Sox2*, *Klf4*, and *c-Myc*. KdVS3 hiPSC line was generated using episomal reprogramming, with the same four reprogramming factors.

*Neuron differentiation*

In the first day of differentiation (or days *in vitro* (DIV)), medium was changed to DMEM/F12 (Gibco, #11320-074) supplemented with 0.1 µg/ml primocin, 4 µg/mL doxycycline, 1% N-2 supplement (Gibco, #17502-048), 1% MEM non-essential amino acid solution (NEAA; Sigma-Aldrich, #M7145), 10 ng/mL human recombinant BDNF (Promokine, #C66212), 10 ng/mL human recombinant NT-3 (Promokine, #C66425), and 0.2 µg/ml Mouse Laminin (Sigma-Aldrich, L2020-1MG). On DIV2, 10,000 cells/well of freshly prepared primary cortical rat astrocytes (isolated as previously described [2]) were added to the neuronal cultures. In DIV3, 2 µM Cytosine β-D-arabinofuranoside (Ara-C; Sigma-Aldrich, C1768-100MG) were added to remove any proliferating cells. From DIV3 onwards, Neurobasal medium (Gibco, #21103-049), supplemented with 20 µg/mL B-27 (Gibco, #0080085SA), 1% GlutaMAX (Gibco, #35050038), 0.1 µg/mL primocin, 4 µg/mL doxycycline, 10 ng/mL human recombinant NT-3, and 10 ng/mL human recombinant BDNF was used. From DIV6 onwards, half of the medium was refreshed three times a week. From DIV10 onwards, medium was supplemented with 2.5% fetal bovine serum (Sigma-Aldrich) to support astrocyte viability. Doxycyline was only added up to and including DIV13. Neuronal cultures were kept at 37°C/5% $CO_2$ during the full differentiation process. For drug testing, compounds in a final concentration of 10 µM and vehicle-controls were added from DIV6, as described in the drug treatment section.

*PCR and sanger sequencing*

Primers targeting *KANSL1* were designed using Primer3Plus [3] and the NCBI Primer-Blast tool. DNA was isolated from hiPSCs by lysing cells with TE buffer (10 mM Tris, 0.1 mM EDTA), 10% NP-40 (Simga-Aldrich,

I8896-50ml), and 10 mg/mL Proteinase K (Thermo Fisher Scientific, E00491). Samples were heated at 56°C for 120 min and at 90°C for 30 min. RNA was isolated from KdVS4 hiPSCs with the Quick-RNA Microprep kit (Zymo Research, R1051) and cDNA was generated by the iScript cDNA synthesis kit (BIO-RAD, 1708891), according to manufacturer's instructions. For both RNA and cDNA samples, PCR was performed using 3.75 µL AmpliTaq Gold 360 master mix (Thermo Fisher Scientific, 4398886) and primers at a final concentration of 0.4 µM. Samples were amplified using the following program: 95°C for 10 min, 35 cycles of 95°C for 30 sec, 58°C for 30 sec, and 72°C for 60 sec, and a final extension at 72°C for 7 min. For H1/H2 haplotyping of all cell lines, gel electrophoresis was performed on the amplicons. For genotyping and identification of *KANSL1* isoforms in KdVS4, amplicons were purified using FastAP (Thermo Fisher Scientific, FEREF0652), Exonuclease I (NEB, M0293), and 1 M/pH 8.0 TrisHCl, by heating samples at 37°C for 15 min and at 80°C for 15 min. Then, sanger sequencing was performed.

Primers: Primers to target intron 12 of *KANSL1* for genotyping of KdVS4 consisted of forward primer 5'-CCAGTTTTAGCCTGCTTGCC-3' and reverse primer 5'- GCATCACAAAGCCACTGTCC-3'. Primers to target exon 10 to exon 14 of *KANSL1* to identify isoforms of *KANSL1* on cDNA (RNA) level consisted of forward primer 5'- CTCACAGGTTACAGCCAGCA-3' and reverse primer 5'- CTGGTGTGGAACAACGGGTA-3'. Primers to target *MAPT* to identify the H1/H2 haplotypes in chromosome 17 consisted of forward primer 5'-GGAAGACGTTCTCACTGATCTG-3' and reverse primer 5'- AGGAGTCTGGCTTCAGTCTCTC-3', as described previously [4].

*Rat astrocytes*

The rodent astrocytes used in this study were derived from embryonic E18 rat brains, as previously described [2]. Animal experiments were conducted in conformity with the Animal Care Committee of the Radboud University Nijmegen Medical Centre, The Netherlands, and conform to the guidelines of the Dutch Council for Animal Care and the European Communities Council Directive 2010/63/EU.

*RNA sequencing*

Library preparation: The library preparation procedure was adapted from a single-cell RNA-seq protocol [5]. The library preparation for RNA samples of hiPSCs from C3, *KANSL1*-CRISPR, and KdVS1 was performed separately. For each sample, 10 ng total RNA (in 0.65 µL) was mixed with 0.1 µL dNTP mix (10 mM each; Invitrogen, 10297018), 0.3 µL nuclease-free water (NF H2O) and 0.4 µL anchored oligo-dT (2.5 µM)

Primer (5'-ACGACGCTCTTCCGATCTNNNNNNNN[10bpindex]TTTTTTTTTTTTTTTTTTTTTTTTTTTTTVN-3', where "N" is any base and "V" is either "A", "C" or "G"; IDT) in a tube containing 7 µL Vapor-Lock

(Qiagen, 981611). Each sample was incubated for 5 min at 65°C and directly placed on ice. Then, first strand reaction mix was added, consisting of 0.4 µL Maxima RT buffer (5X) (Thermo Fisher Scientific, EP0751), 0.05 µL RNasin Plus (Promega, N2611) and 0.1 µL Maxima H Minus Reverse Transcriptase (Thermo Fisher Scientific, EP0751). Reverse transcription was performed by incubation at 50°C for 30 min, and terminated by heating at 85°C for 5 min. 2 µL RT product was mixed with 7.7 µL NF H2O, 2.5 µL Second Strand Buffer (Invitrogen, 10812014), 0.25 µL dNTP mix (10 mM each), 0.35 µL DNA polymerase I (E.coli; NEB, M0209), 0.09 µL DNA ligase (E. coli; NEB, M0205) and 0.09 µL Ambion RNase H (E. coli; Invitrogen, AM2293). Second strand synthesis was performed by incubation at 16°C for 150 min, followed by 75°C for 20 min. 0.5 µL Exonuclease I (NEB, M0293) were added per sample and incubated at 37°C for 60 min. cDNA samples were pooled per sets of 8-9, randomized across three pools. Vapor-Lock was removed, and samples were added up with NF H2O to a total volume of 121.1 µL. Each pool of samples was purified using 95 µL beads buffer (20% PEG-8000 in 2.5 M NaCL, final concentrations) and 50 µL Ampure XP Beads (Beckman Coulter, A63881), and eluted in 7 µL NF H2O.

Tagmentation was performed per pool by adding 3.0 µL double-stranded cDNA sample to 5.0 µL Nextera TD buffer (Illumina, 15027866), 2.5 µL NF H2O and 1.0 µL TDE1 Enzyme (Illumina, 15027865), incubated at 55°C for 5 min. Samples were directly placed on ice for at least 3 min. The reaction was terminated by adding 12 µL Buffer PB (QiaQuick, 19066) and incubating for 5 min at room temperature. Samples were purified using 48 µL Ampure XP beads and eluted in 10 µL NF H2O. Next, each sample was mixed with 2 µL P5 primer (10 µM), (5'-AATGATACGGCGACCACCGAGATCTACAC[i5]ACACTCTTTCCCTACACGACGCTCTTCCGATCT-3'; IDT), 2 µL P7 primer (10 µM) (5'-CAAGCAGAAGACGGCATACGAGAT[i7]GTCTCGTGGGCTCGG-3'; IDT), 6 µL NF H2O, and 20 µL NEBNext High-Fidelity 2X PCR Master Mix (NEB, M0541). Amplification was performed using the following program: 72°C for 5 min, 98°C for 30 sec, 15 cycles of: 98°C for 10 sec, 66°C for 30 sec, 72°C for 1 min, and a final step at 72°C for 5 min. Next, samples were purified using 32 µL Ampure XP beads and eluted in 12 µL NF H2O. Libraries were visualized by electrophoresis on a 1% agarose and 1X TAE gel containing 0.3 µg/mL ethidium bromide (Invitrogen, 15585011). Gel extraction was performed to select products between 200 – 1000 bp using the Nucleospin Gel and PCR Clean-up kit (Macherey-Nagel, 740609). Samples were eluted in 11 µL NF H2O. cDNA concentration was measured by Qubit dsDNA HS Assay kit (Invitrogen, Q32854). Product size distributions were visualized using Agilent's Tapestation system (D5000 ScreenTape and Reagents, 5067-5588/9).

*RNA-seq data pre-processing*

Base calls were converted to fastq format and demultiplexed using Illumina's bcl2fastq conversion software v2.16.0.10 tolerating one mismatch per library barcode. Reads were filtered for a valid unique molecular identifier (UMI) and sample barcode, tolerating one mismatch per barcode. Trimming was performed using Trimmomatic v0.33 [6]. For hiPSCs, trimmed reads were mapped to the human reference genome (GRCh38.p12; release 29, Ensembl 94), whereas for hiPSC-derived neurons co-cultured with rat astrocytes trimmed reads were mapped to a combined human GRCh38.p12 and rat reference genome (Rnor_6.0). Mapping was performed using STAR v2.5.1b [7], with default settings. Uniquely mapped reads (mapping quality of 255) were extracted and read duplicates were removed using the UMI-tools software package [8]. The reads in BAM files were count with HTSeq v0.9.1[9], using GRCh38.p12 reference transcriptome for hiPSCs, and the combined GRCh38.p12 and Rnor_6.0 reference transcriptome for the hiPSC-derived neuronal co-cultures, to separate counts from human neurons and rat astrocytes.

*RNA-seq data analysis for drug screening*

Only transcripts with an overall cpm > 10 were considered, determined by *filterByExpr* of EdgeR, resulting in 8817 included for DE analysis. Counts were voom-transformed with limma. A linear regression model was fit, in which the voom-transformed expression values were modelled as a function of the condition. Each KdVS line was compared to its MEA-batch control(s); therefore, the gene expression signature of the *KANSL1*-CRISPR was compared to C3, KdVS1 was compared to C3, and both KdVS2 and KdVS3 were compared to C1 and C2. Genes with a BH-corrected $p<0.05$ were considered as differentially expressed.

*CLCN4 RNA interference*

<u>HEK293T cells culturing:</u> HEK293T cells (ATCC, CRL-3216) were cultured in high-glucose DMEM (Sigma, D0819) supplemented with 10% Sodium-Pyruvate (Sigma, S8636), 1% Pen/strep and 10% FCS in 10 cm petridishes. Cells were split using 0.05% Trypsin-EDTA (GIBCO, 25300054) in 1:10 dilution for the generation of lentiviral particles, using four plates per construct.

<u>Lentiviral particles preparation:</u> Lentiviruses were generated by co-transfecting each lentiviral vector, the psPAX2 packaging vector (Addgene, #12260) and the VSVG envelope glycoprotein vector pMD2-G (Addgene, #12259) into HEK293T cells, using calcium phosphate precipitation. The supernatant of culture medium was collected 48 hours after transfection and filtered through a 0.45 µm syringe filter. Lenti-X Concentrator (Takara, 631231) was added in a 1:4 ratio to the filtered medium, which was incubated overnight at 4°C. After centrifugation, the supernatant was removed and the cell pellet was dissolved in PBS. Viral particles were stored at -80°C.

*Quantification of mRNA by RT-qPCR*

Primers targeting *CLCN4*: Forward primer 5’-TGGTAGTGTCCTCCGGTCTG-3’; reverse primer 5’-AAAGGCAACAGAGACTCCAGC-3’.

RT-qPCR program: Initial denaturation step at 95°C for 10 min, followed by 40 cycles of 95°C for 15 sec and 60°C for 30 sec, followed by generation of a melting curve.

*Immunocytochemistry*

After fixation, hiPSCs-derived neurons were washed 3x with 1X PBS and permeabilized using 0.2% triton X-100 (Sigma-Aldrich, #9002-93-1). Cells were washed 3x with 1X PBS and incubated with blocking buffer, consisting of 1X PBS, 5% normal horse serum (Gibco, 26050070), 5% normal goat serum (Invitrogen, 10000C), 5% normal donkey serum (Jackson Immuno Research, 017-000-121), 0.1% bovine serum albumin (Sigma-Aldrich, A9418-10G), 1% glycine, 0.4% triton X-100, and 0.1% lysine (Sigma-Aldrich, P4707) for 1 hour at room temperature. Primary antibodies were diluted in blocking buffer and were incubated overnight at 4°C. Cells were washed 3x with 1X PBS and incubated with secondary antibodies for 1 hour at room temperature. Cells were washed 3x with 1X PBS and Hoechst (Thermo Fisher Scientific, #H3570) was incubated for 10 minutes at room temperature (1:10,000 in 1X PBS). Cells were washed one time with 1X PBS, and coverslips were embedded in mounting medium (DAKO, #S3023). Primary antibodies that were used: guinea pig anti-MAP2 (Synaptic Systems, #188044, 1:1000), rabbit anti-CLCN4 (Sigma-Aldrich, HPA063637, 1:500), mouse anti-MAP2 (Abcam, ab11267, 1:1000), and rabbit anti-GFAP (Abcam, ab7260, 1:1000). Secondary antibodies that were used: goat anti-rabbit Alexa fluor 647 (Molecular Probes, A21245, 1:500), goat anti-guinea pig Alexa fluor 568 (Invitrogen, A11075, 1:500), and goat anti-mouse Alexa fluor 488 (Invitrogen, A11029, 1:1000).

Querying the LINCS database

Connectivity score calculation: This quantifies the extent to which upregulated query genes appear towards the top of the ordered perturbational gene expression profile, and downregulated genes towards the bottom of the ordered perturbational gene expression profile. For a given query q and a gene expression profile r, the weighted connectivity score is calculated as

$$WCS_{q,r} = \begin{cases} KSup - KSdown, & if\ sign(KSup) = sign(KSdown) \\ 0, & otherwise \end{cases} \quad (1)$$

Where KS is the Kolmogorov-Smirnov statistic, calculated as

$$KS = \begin{cases} a = \max\limits_{1 \le j \le t} \left[ \frac{j}{t} - \frac{V(j)}{n} \right], & if\ a > b \\ -b = -\max\limits_{1 \le j \le t} \left[ \frac{V(j)}{n} - \frac{(j-1)}{t} \right], & otherwise \end{cases} \tag{2}$$

KS is divided into KSup and KSdown, referring to up- and downregulated genes. Let n be the total number of genes in an ordered gene expression profile, and t the number of up- or downregulated query genes. *V(j)* is the position of any gene j in the ordered gene expression profile. Gene *j* is any gene in the up- or downregulated query gene list, where *j* = 1, 2, …, *t*. Subsequently, weighted connectivity scores are normalized to allow comparison of connectivity scores across cell types and perturbation types. Given a vector *w* of weighted connectivity scores profiled in cell line *c* and perturbagen type *t*, the normalized connectivity score is calculated as

$$NCS_{c,t} = \begin{cases} w_{c,t} \ / \ \mu^{+}_{c,t}, & if\ sign(w_{c,t}) > 0 \\ w_{c,t} \ / \ \mu^{-}_{c,t}, & otherwise \end{cases} \tag{3}$$

Where $\mu^{+}_{c,t}$ is the mean of all positive weighted connectivity scores in *w*, and $\mu^{-}_{c,t}$ is the mean of all negative weighted connectivity scores in *w*. The normalized connectivity score $NCS_{c,\ t}$ is then adjusted to retain the sign of $W_{c,\ t}$.

Finally, since only a subset of compounds has been tested on a neuronal cell type by the LINCS consortium, we queried our KdVS signatures against all signatures of a compound combined. We calculated a compound-centric measure of connectivity to the query genes, summarized across cell lines. We will further refer to this as the cell-summarized score ($Q_{c,t}$), i.e. the quantile score. Let $ncs_p$ be a vector of normalized connectivity scores for a specific perturbagen *p* relative to query *q* across all cell lines where *p* was profiled. To mitigate the effect of outliers in this procedure, we only considered compounds with at least 10 gene expression profiles in the database that showed connectivity to the query (e.g. the weighted connectivity score differs from zero). $Q_{high}$ and $Q_{low}$ are then the upper and lower quantiles of this vector, respectively. Following the LINCS consortium, we used $Q_{low}$ = 33 and $Q_{high}$ = 67. The cell-summarized score was then calculated as

$$Q_{c,t} = \begin{cases} Q_{high}(ncs_p), & if\ |Q_{high}(ncs_p)| \ge |Q_{low}(ncs_p)| \\ Q_{low}(ncs_p), & otherwise \end{cases} \tag{4}$$

*Data visualization*

Data visualization was performed in R. Plots showing MEA data and synapsin puncta over time depict the mean ± standard error of the mean (SEM) per condition. Spider charts were generated using *radarchart* function of fmsb R package. Heatmaps were generated using *pheatmap*, with hierarchical clustering performed on rows and columns, if applicable. Venn diagrams were generated using ggVennDiagram [10]. Correlation plots were generated using *ggcorrplot*. Pie charts were generated using *pie* function of graphics. PCA plots and all other remaining figures were made using ggplot2.

### Supplementary figures

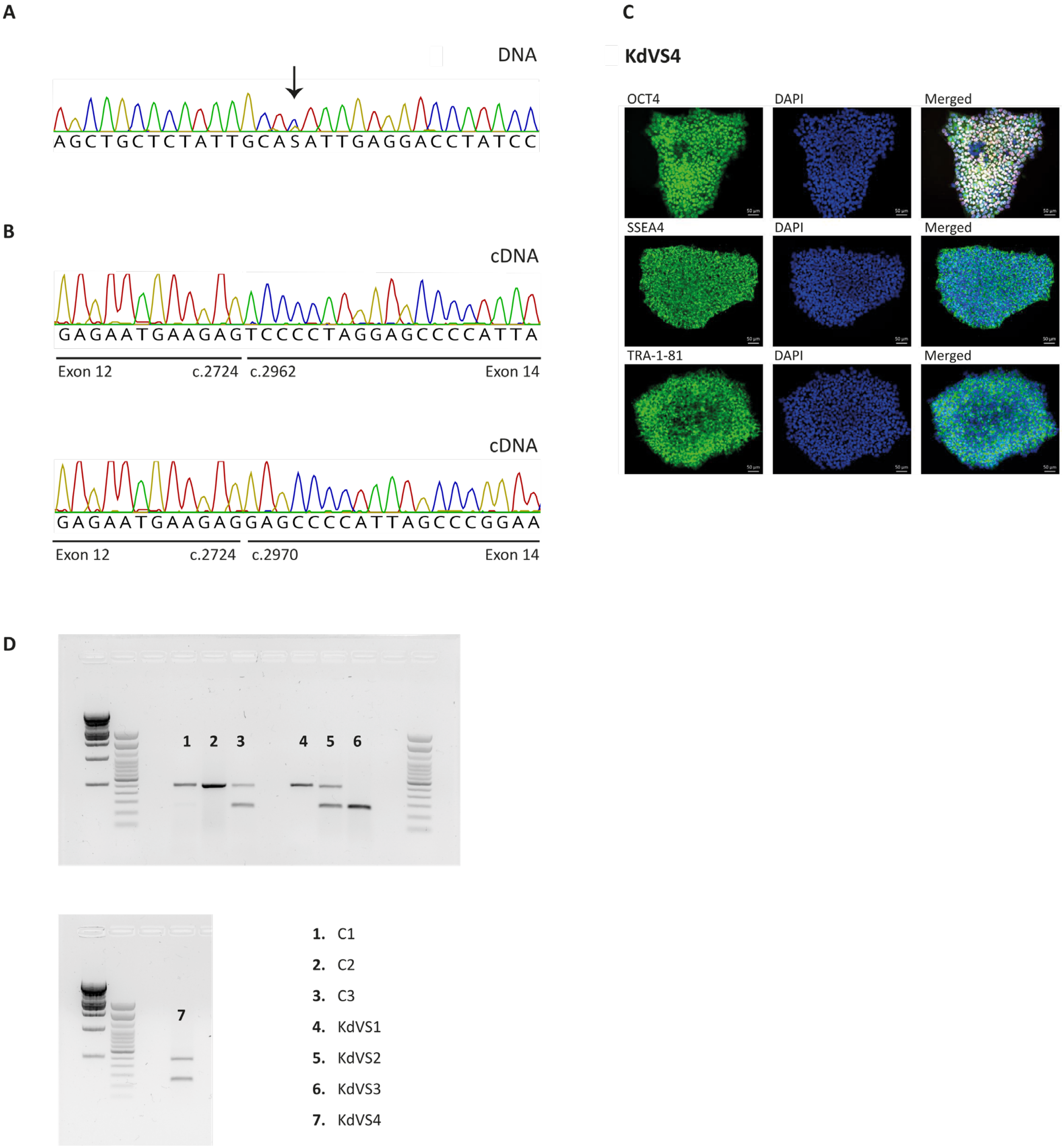


**Figure S1. hiPSCs characterization.**

**(A)** Sanger sequencing chromatogram depicting the DNA sequence of *KANSL1* in KdVS4 hiPSCs. The arrow points to a C>G substitution at position chr17:g.44110559, encoding for the last base pair of intron 12 in *KANSL1* (c.2725-1G>C). **(B)** Sanger sequencing chromatograms showing the cDNA sequence generated from RNA isolated from KdVS4 hiPSCs. The chromatograms revealed two isoforms, in which 238 bp (top panel) and 246 bp (bottom panel) are skipped. In both isoforms, exon 13 and a part of exon 14 is missing. **(C)** Immunostaining images of KdVS4 hiPSCs, showing expression of pluripotency markers OCT4, SSEA4, and TRA-1-81 (scale bar = 50 µm). **(D)** Gel electrophoresis performed on DNA samples from KdVS and control lines. Bands represent different haplotypes in chromosome 17, with the top band indicative of a H1 haplotype and the bottom band indicative of a H2 haplotype.

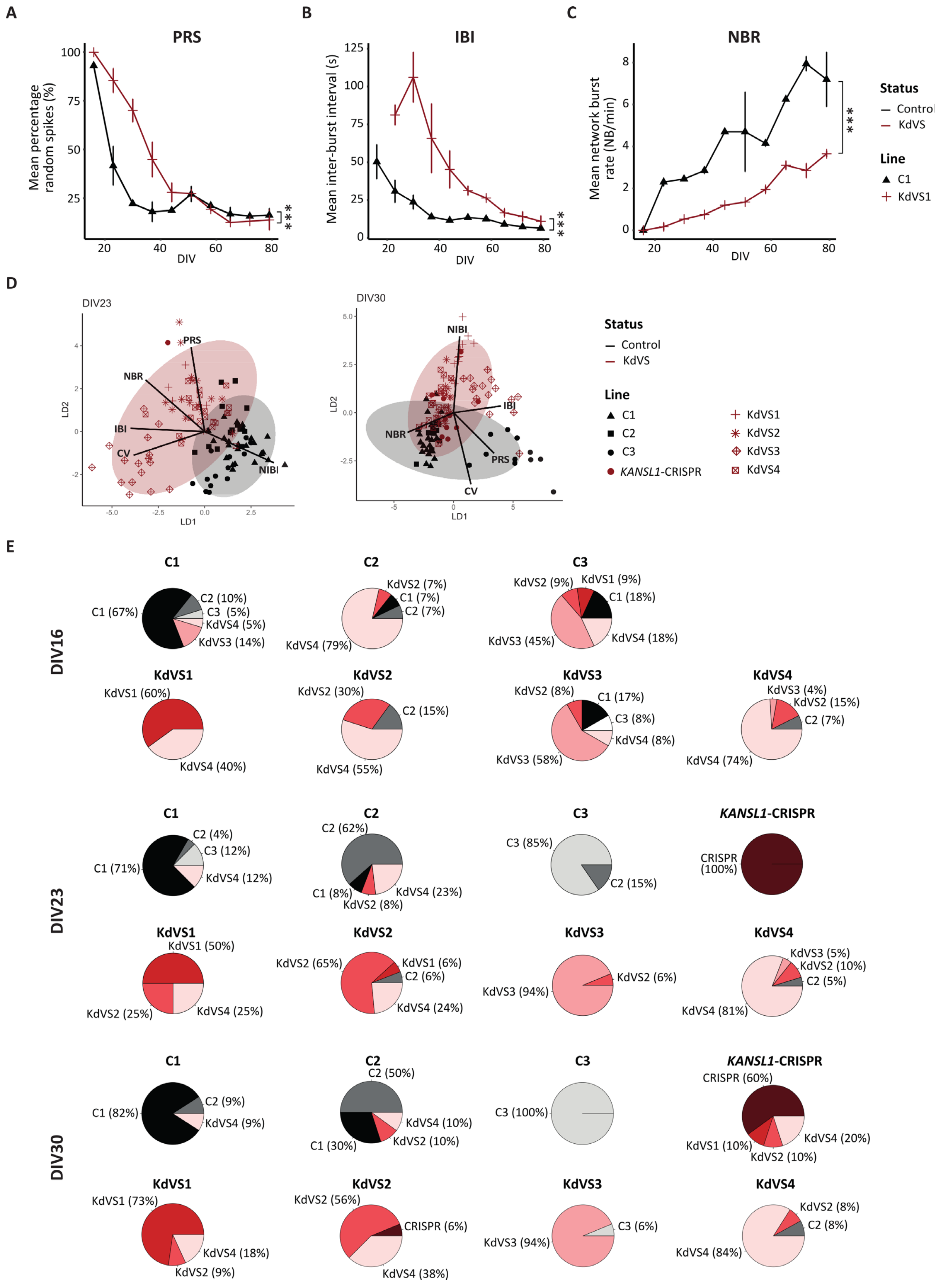
A
PRS
Mean percentage random spikes (%)
DIV
***
B
IBI
Mean inter-burst interval (s)
DIV
***
C
NBR
Mean network burst rate (NB/min)
DIV
***
Status
Control
KdVS
Line
C1
KdVS1
D
DIV23
DIV30
LD1
LD2
PRS
NBR
IBI
CV
NIBI
Status
Control
KdVS
Line
C1
C2
C3
KANSL1-CRISPR
KdVS1
KdVS2
KdVS3
KdVS4
E
DIV16
DIV23
DIV30
C1
C2
C3
KdVS1
KdVS2
KdVS3
KdVS4
KANSL1-CRISPR

**Figure S2. Supporting data on network activity recordings in KdVS and control hiPSC-derived neurons.** **(A-C)** Plots showing parameters describing neuronal network activity in KdVS1 ($n_{wells}$=3) and control C1 neurons ($n_{wells}$=2), consisting of the PRS **(A)**, the IBI **(B)**, and the NBR **(C)**. Data represent means ± SEM, and are shown for DIV16-79. Significance was determined by one-way ANOVA followed by a Tukey HSD post-hoc test. *** *p*-value<0.001, ** *p*-value<0.01, * *p*-value<0.05. **(D)** Linear discriminant analysis (LDA) performed on 5 MEA parameters affected in KdVS (PRS, IBI, NBR, NIBI, $CV_{NIBI}$), for KdVS (red) and controls neurons (black), at DIV23 (left panel) and DIV30 (right panel). The arrows indicate the direction of MEA parameters in KdVS. Shapes represent different cell lines. **(E)** Pie charts showing the predicted membership for each cell line based on LDA on all 10 MEA parameters describing the neuronal network activity, for DIV16, DIV23, and DIV30. $CV_{NIBI}$ = coefficient of variation calculated on the NIBI ; DIV = days *in vitro*; MEA = microelectrode array; PRS = percentage of random spikes; IBI = inter-burst interval; NBR = network burst rate; NIBI = NB inter-burst interval.

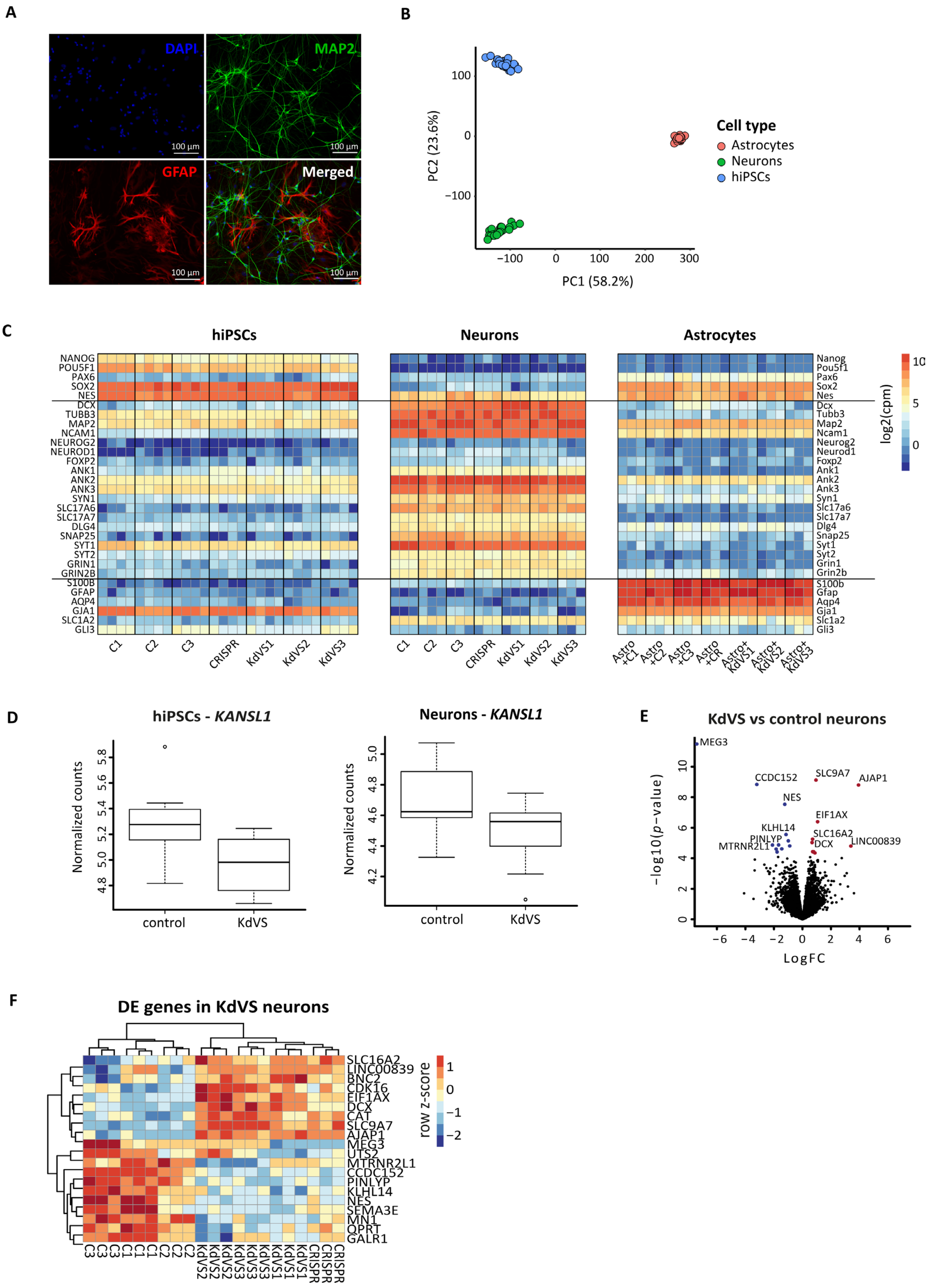

A
DAPI
MAP2
GFAP
Merged
100 µm
B
Cell type
Astrocytes
Neurons
hiPSCs
PC2 (23.6%)
PC1 (58.2%)
C
hiPSCs
Neurons
Astrocytes
NANOG
POU5F1
PAX6
SOX2
NES
DCX
TUBB3
MAP2
NCAM1
NEUROG2
NEUROD1
FOXP2
ANK1
ANK2
ANK3
SYN1
SLC17A6
SLC17A7
DLG4
SNAP25
SYT1
SYT2
GRIN1
GRIN2B
S100B
GFAP
AQP4
GJA1
SLC1A2
GLI3
C1
C2
C3
CRISPR
KdVS1
KdVS2
KdVS3
log2(cpm)
D
hiPSCs - KANSL1
Neurons - KANSL1
Normalized counts
control
KdVS
E
KdVS vs control neurons
MEG3
CCDC152
SLC9A7
AJAP1
NES
EIF1AX
KLHL14
PINLYP
MTRNR2L1
SLC16A2
DCX
LINC00839
−log10(p−value)
LogFC
F
DE genes in KdVS neurons
SLC16A2
LINC00839
BNC2
CDK16
EIF1AX
DCX
CAT
SLC9A7
AJAP1
MEG3
UTS2
MTRNR2L1
CCDC152
PINLYP
KLHL14
NES
SEMA3E
MN1
QPRT
GALR1
row z-score

**Figure S3. Supporting data on the gene expression profiles of KdVS hiPSCs and hiPSC-derived neurons.** **(A)** hiPSC-derived neurons co-cultured with rat astrocytes (DIV23) were stained for DAPI, MAP2, and GFAP. **(B)** Principal component analysis (PCA) on gene expression profiles of hiPSCs, hiPSC-derived neurons and rat astrocytes, considering all genes for which a human homologue was available. PC1 and PC2 are shown. Samples are colored per cell type, showing that cell type explains the largest variation of the transcriptomic profiles. **(C)** Heatmap depicting gene expression level of stem cell/ neural progenitor cell (top section), neuronal and synaptic (middle section), and glial marker genes (bottom section) in hiPSCs and hiPSC-derived neurons, confirming the cell identity of each sample. Left Y axis indicates human gene symbols, while right Y axis indicates the rat homologs. Voom-transformed and batch-corrected counts per million (log2 scale) are shown. **(D)** Boxplots showing normalized *KANSL1* expression in hiPSCs and hiPSC-derived of KdVS and controls. **(E)** Volcano plot showing DE genes in KdVS hiPSC-derived neurons (DIV30) compared to control neurons. **(F)** Heatmap showing expression of neuronal DE genes in KdVS ($n_{wells}$=12) and control ($n_{wells}$=9) neurons.

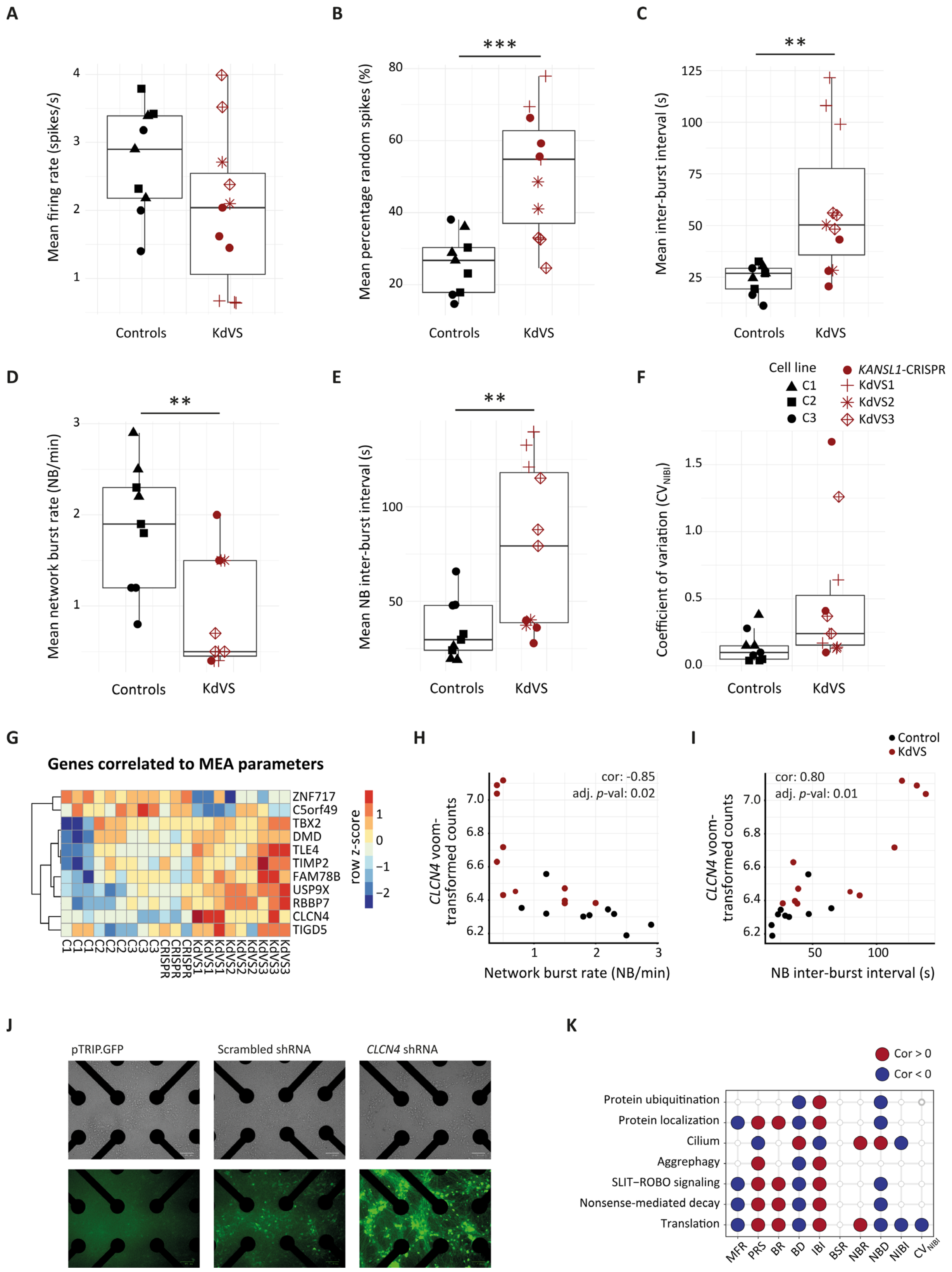
A
B
C
Mean firing rate (spikes/s)
Mean percentage random spikes (%)
Mean inter-burst interval (s)
Controls
KdVS
D
E
F
Mean network burst rate (NB/min)
Mean NB inter-burst interval (s)
Coefficient of variation (CV_NIBI)
Cell line
C1
C2
C3
KANSL1-CRISPR
KdVS1
KdVS2
KdVS3
G
Genes correlated to MEA parameters
ZNF717
C5orf49
TBX2
DMD
TLE4
TIMP2
FAM78B
USP9X
RBBP7
CLCN4
TIGD5
row z-score
H
I
Control
KdVS
cor: -0.85
adj. p-val: 0.02
cor: 0.80
adj. p-val: 0.01
CLCN4 voom-transformed counts
Network burst rate (NB/min)
NB inter-burst interval (s)
J
pTRIP.GFP
Scrambled shRNA
CLCN4 shRNA
K
Cor > 0
Cor < 0
Protein ubiquitination
Protein localization
Cilium
Aggrephagy
SLIT–ROBO signaling
Nonsense-mediated decay
Translation
MFR
PRS
BR
BD
IBI
BSR
NBR
NBD
NIBI
CV_NIBI

**Figure S4. Supporting data on gene expression and neuronal network activity correlations.**

**(A-F)** Boxplots showing the MFR **(A)**, PRS **(B)**, IBI **(C)**, NBR **(D)**, NIBI **(E)**, and the $CV_{NIBI}$ **(F)** of hiPSC-derived neurons (DIV30) included in the MEA-seq experiments (KdVS $n_{wells}$=11, controls $n_{wells}$=9). Shapes represent different cell lines. Data represent mean ± SEM. Significance was determined by a one-sample *t*-test, comparing KdVS to controls. **(G)** Heatmap showing expression of the genes that significantly correlated with at least one MEA parameter and that were also differentially expressed in KdVS neurons (KdVS $n_{wells}$=12, control $n_{wells}$=9; $p<0.05$). Voom-transformed and batch-corrected counts per million (log2 scale) were scaled per gene (row z-score). **(H-I)** Scatterplots showing expression of *CLCN4* (y-axis) against the NBR **(H)** and NIBI **(I)**, in all samples included in MEA-seq. **(J)** Images of KdVS1 neuronal cell density (top panels) and GFP expression (bottom panels) after transduction with lentivirus expressing the empty vector pTRIP.GFP, the scrambled shRNA, or the *CLCN4*-targeting shRNA. **(K)** Correlation plot showing gene sets (Gene Ontology (GO) terms and Reactome pathways) significantly enriched (adj. $p<0.05$) for genes that either positively (red) or negatively correlated with the MEA parameters. ****p*-value<0.001, ** *p*-value<0.01, * *p*-value<0.05. $CV_{NIBI}$ = coefficient of variation calculated on the NIBI; IBI = inter-burst interval; MFR = mean firing rate; NBR = network burst rate; NIBI = NB inter-burst interval; PRS = percentage of random spikes.

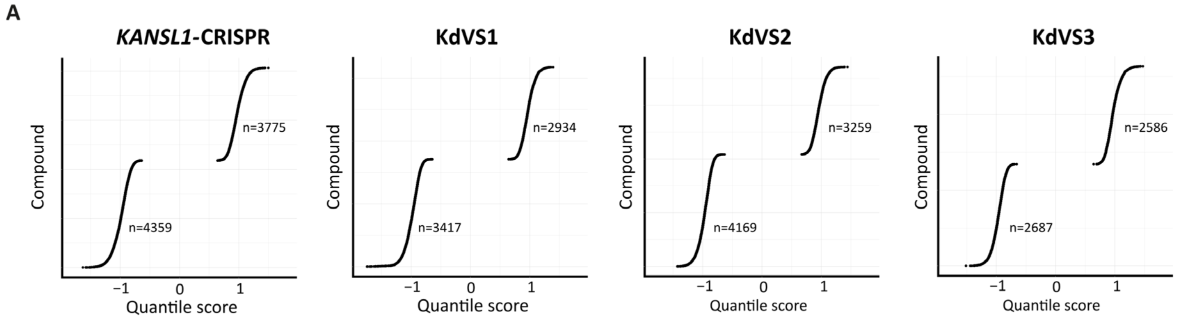


**Figure S5. Supporting data on connectivity scores determined between query genes of individual KdVS signatures and perturbation signatures from the LINCS database.**

**(A)** Scatter plot showing quantile scores ($Q_{c,t}$) for all compounds with overall positive ($Q_{c,t} > 0$) or negative ($Q_{c,t} < 0$) connectivity, determined using query genes per individual KdVS line.

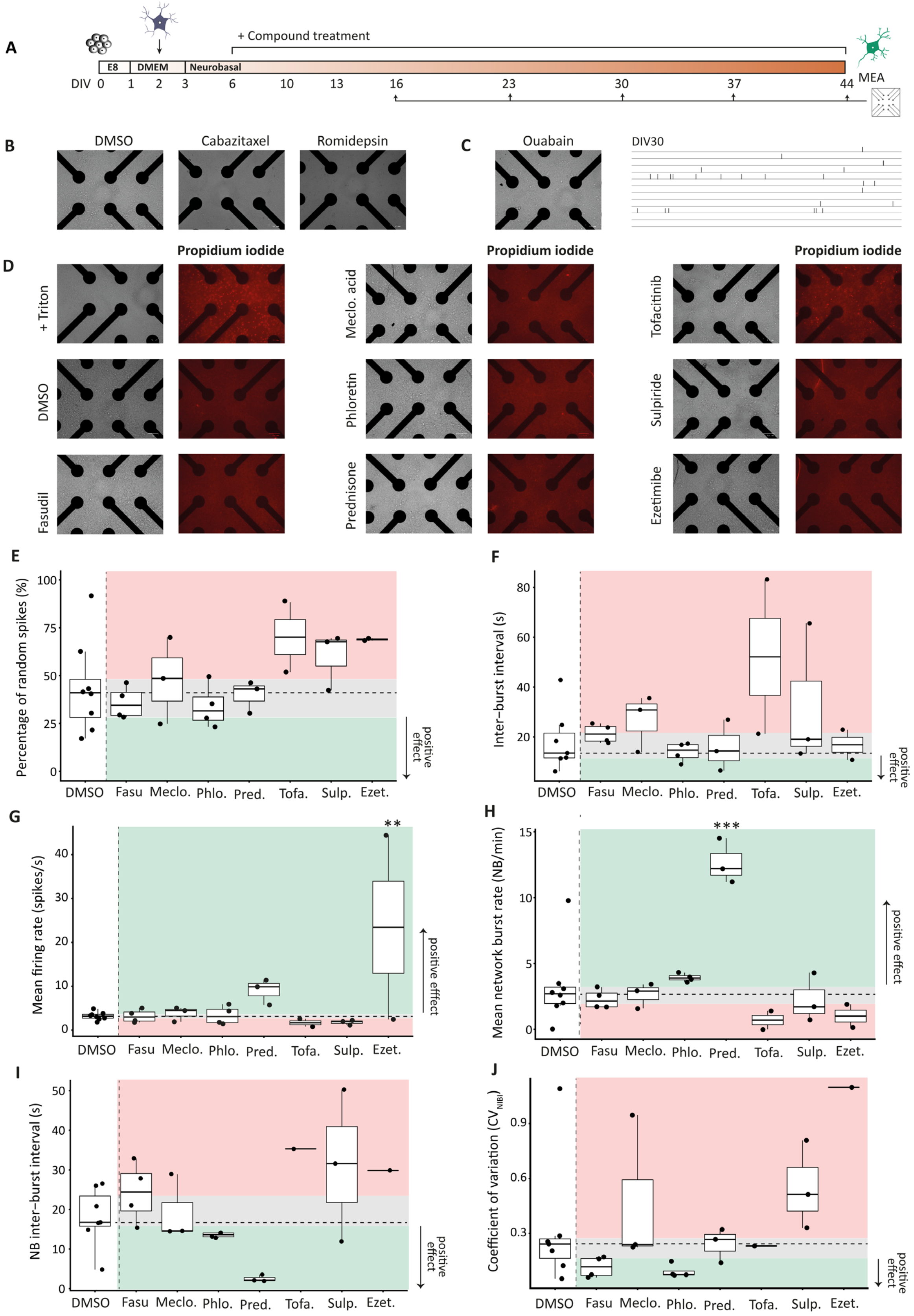

A
+ Compound treatment
E8
DMEM
Neurobasal
DIV 0 1 2 3 6 10 13 16 23 30 37 44
MEA
B
DMSO
Cabazitaxel
Romidepsin
C
Ouabain
DIV30
D
Propidium iodide
+ Triton
DMSO
Fasudil
Meclo. acid
Phloretin
Prednisone
Tofacitinib
Sulpiride
Ezetimibe
E
Percentage of random spikes (%)
positive effect
F
Inter−burst interval (s)
G
Mean firing rate (spikes/s)
**
H
Mean network burst rate (NB/min)
***
I
NB inter-burst interval (s)
J
Coefficient of variation ($CV_{NIBI}$)
DMSO
Fasu
Meclo.
Phlo.
Pred.
Tofa.
Sulp.
Ezet.

**Figure S6. Supporting data on effect of compounds on *KANSL1*-CRISPR hiPSC-derived neurons**.

**(A)** Experimental workflow. At DIV0, *KANSL1*-CRISPR hiPSCs were plated on MEA and differentiated into neurons up to DIV44. At DIV2, rat astrocytes were added to support neuronal maturation. From DIV6 onwards, the compound was added in the medium, while changing the medium every other day to maintain a final compound concentration of 10 µM. Neuronal network activity was measured at DIV16, 23, 30, 37 and 44. **(B)** Cell density images of *KANSL1*-CRISPR derived neurons treated either with DMSO (0.1%), 10 µM of cabazitaxel or romidepsin. Cabazitaxel and romidepsin induced cell death and detachment. **(C)** Cell density image and raster plot showing neuronal network activity in ouabain-treated cultures (10 µM). Even though average cell density, network activity was almost null (DIV30). **(D)** Cell density images of *KANSL1*-CRISPR derived neurons treated with DMSO (0.1%) or 10 µM of compound. On DIV44, cells were treated with propidium iodide (in red) to evaluate cell death. One well was treated with triton and propidium iodide, as a positive control. **(E-J)** Effect of the compounds ($n_{wells}$=2-4) and DMSO ($n_{wells}$=8) on different activity parameters. The median value for DMSO-treated *KANSL1*-CRISPR neurons is indicated with a horizontal dashed line, and the Q1-Q3 range is highlighted in grey. The expected range for the effect of compounds is indicated in green (positive effect, towards the control level). Measurements in the red area are indicative of adverse effect of the compound (negative effect, reinforced KdVS phenotype). Difference between groups was tested with one-way ANOVA, followed by a Dunnett's post-hoc test comparing all compounds to DMSO-treated *KANSL1*-CRISPR neurons. * adj. *p*-value <0.05; ** <0.01; *** <0.001, as respect to DMSO-treated *KANSL1*-CRISPR neurons. Fasu = fasudil, Meclo = meclofenamic acid, Phlo = phloretin, Pred = prednisone, Tofa = tofacitinib, Sulp = sulpiride, Ezet = ezetimibe, BR = burst rate; $CV_{NIBI}$ = coefficient of variation calculated on the NIBI; IBI = inter-burst interval; NBR = network burst rate; NIBI = NB inter-burst interval; MFR = mean firing rate; PRS = percentage of random spikes.

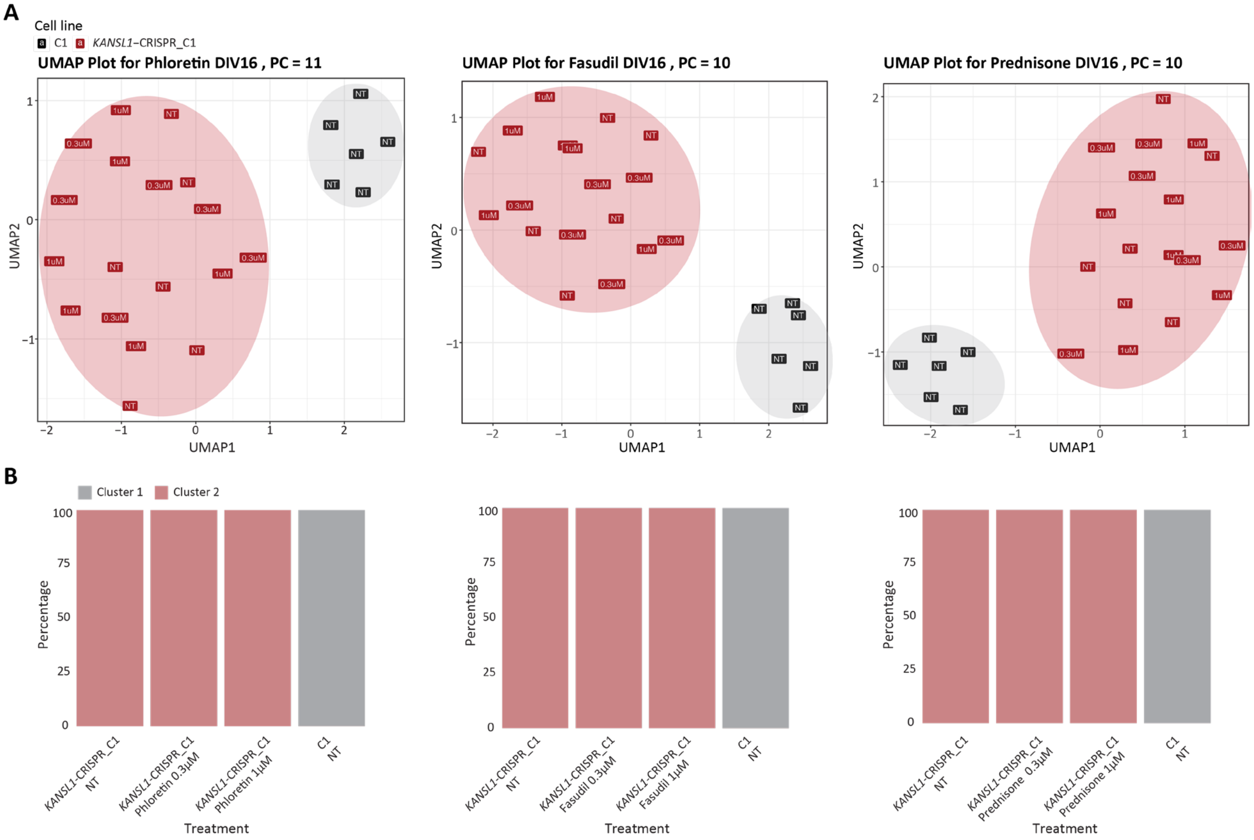


**Figure S7**. **Supporting data on the effect of fasudil, phloretin and prednisone on neuronal network activity in *KANSL1*-CRISPR_C1-derived neurons.**

**(A)** UMAPs were generated from principal components (PCs) obtained from network activity parameters for C1 and *KANSL1*-CRISPR_C1-derived neurons (DIV16) treated with phloretin (11 PCs), fasudil (10 PCs), and prednisone (10 PCs). Compounds were added at final concentrations of 0.3 µM and 1 µM. Hierarchical clustering analysis was then performed on the sample distances based on the two UMAP dimensions. The ellipses group the recordings assigned to each cluster. **(B)** Proportion of recordings assigned to each hierarchical cluster per condition.

**Supplementary tables**

**Table 1.** Differentially expressed genes between KdVS and control hiPSC-neurons.

**Table 2.** MEA-seq corrlations and GSEA.

**Table 3.** Differentially expressed genes between KdVS and control hiPSC-neurons, within batches, for computational drug screening.

**Table 4.** Overview of top 10 selected compounds with negative connectivity with KdVS signatures**.**

| Compound | Mechanism | Selection criteria | FDA approved | Concentration | Manufacturer |
|---|---|---|---|---|---|
| Cabazitaxel | Microtubule inhibitor | MOA and TNQ (KdVS all) | X | 10 mM in DMSO | Bio-connect, HY-15459 |
| Ezetimibe | Cholesterol inhibitor | TNQ (KANSL1) | X | 10 mM in DMSO | Bio-connect, HY-17376_10mM |
| Meclofenamic acid | Cyclooxyygenase inhibitor | TNQ (KANSL1) | | 1 mg as powder | Sigma-Aldrich, M4531-1G |
| Ouabain | ATPase inhibitor | TNQ (KdVS all) | | 10 mM in DMSO | Bio-connect, S4016_10mM |
| Romidepsin | HDAC inhibitor | MOA | X | 1 mg as powder | Bio-connect, HY-15149_1mg |
| Sulpiride | Dopamine receptor antagonist | TNQ (KANSL1) | | 100 mg as powder | Tocris, 0894 |
| Tofacitinib | JAK inhibitor | TNQ (KANSL1) | X | 10 mM in DMSO | Bio-connect, S5001_10mM |
| Prednisone | Glucocorticoid receptor agonist | MOA | X | 10 mM in DMSO | Bio-connect |
| Phloretin | Sodium/glucose co-transporter inhibitor | TNQ (KdVS all) | | 10 mM in DMSO | Bio-connect |
| Fasudil | Rho-associated kinase inhibitor | TNQ (KdVS all) | | 10 mM in DMSO | Bio-connect |

The top 10 compounds selected for experimental validation in KdVS hiPSC-derived neurons are shown, including their mechanism of action, the selection criteria, manufacturer and original concentration. Selection based on MOA indicates that compounds were selected based on the top overlapping mechanisms of action between compounds with a negative connectivity. Selection based on the top negative quantile indicates compounds with the highest negative quantile scores that overlapped between all KdVS lines (KdVS all), or between the *KANSL1* mutation lines (KANSL1). FDA = U.S. Food and Drug Administration; HDAC = histone deacetylase; JAK = Janus kinase; MOA = mechanism of action. TNQ = Top negative quantile
